# Age-related epithelial defects limit thymic function and regeneration

**DOI:** 10.1101/2021.12.16.472014

**Authors:** Anastasia I. Kousa, Lorenz Jahn, Kelin Zhao, Angel E. Flores, David Granadier, Kirsten Cooper, Julie M. Sheridan, Andri Lemarquis, Lisa Sikkema, Kimon V. Argyropoulos, Jennifer Tsai, Amina Lazrak, Katherine Nichols, Nichole Lee, Romina Ghale, Florent Malard, Hana Andrlova, Antonio L.C. Gomes, Enrico Velardi, Salma Youssef, Marina B. da Silva, Melissa Docampo, Roshan Sharma, Linas Mazutis, Verena C. Wimmer, Kelly L. Rogers, Susan DeWolf, Brianna Gipson, Manu Setty, Dana Pe’er, Nancy R. Manley, Daniel H.D. Gray, Jarrod A. Dudakov, Marcel R.M. van den Brink

## Abstract

The thymus is essential for establishing adaptive immunity yet undergoes age-related atrophy leading to compromised immune responsiveness. The thymus is also extremely sensitive to acute insult and although capable of regeneration, this capacity declines with age. Focusing on non-hematopoietic stromal cells, and using single-cell and spatial transcriptomics, lineage-tracing, and advanced imaging, we discovered two atypical thymic epithelial cell (TEC) states that emerged with age. Age-associated (aa)TECs formed atypical high-density epithelial clusters that were devoid of thymocytes, an accretion of non-functional thymic tissue that worsened with age and exhibited features of partial epithelial-to-mesenchymal transition (EMT). *In silico* interaction analysis revealed that aaTEC emergence drew tonic signals from other TEC populations at baseline, acting as a sink for TEC growth factors. Following damage, aaTEC expanded substantially, further perturbing trophic pathways, and correlating with defective regeneration of the involuted thymus. These findings define a unique feature of thymic involution linked to immune aging.

## INTRODUCTION

Thymic T cell differentiation requires the close interaction between thymocytes and supporting stromal (epithelial, endothelial, and mesenchymal) cells that comprise the thymic microenvironment (*1, 2*). Yet, despite the importance of the thymus for generating a broad T cell repertoire that can repel infection and maintain immunological tolerance, the generation of new T cells declines precipitously due to age-related thymic involution. Thymic involution is a chronic degenerative process that accelerates upon puberty and is characterized by significant atrophy, disrupted stromal architecture, reduced export of new naïve T cells and, ultimately, diminished responsiveness to new antigens (*3–5*). Along with its chronic functional decline with age, the thymus is also extremely sensitive to acute damage caused by infection or common cancer therapies such as cytoreductive chemo- or radiation therapy (*6*). Although the thymus harbors tremendous capacity for endogenous repair, this regenerative capacity also declines with age (*7–9*), such that profound thymic damage caused by the conditioning regimens required for hematopoietic cell transplantation (HCT) lead to prolonged T cell lymphopenia, an important contributor to transplant-related morbidity and mortality due to infections and malignant relapse (*10–12*). In fact, thymic function can be a positive prognostic indicator of HCT outcomes (*13–15*).

The thymic stroma is composed of highly specialized thymic epithelial cells (TECs), endothelial cells (ECs), mesenchymal cells like fibroblasts (FBs), dendritic cells (DCs), innate lymphoid cells (ILCs), and macrophages; all of which are crucial for aspects of T cell differentiation (*1*). Recent advances in single cell technology have provided new insights into the heterogeneity of TEC in young (*16–19*) and aged mice (*20*), and in humans (*21*); how TEC orchestrate T cell differentiation and how dysfunction in these processes is linked to autoimmunity and immunodeficiency. However, perhaps as a consequence of this complexity, the mechanisms underlying thymic involution and regeneration after damage remain poorly understood (*7–9*). Here we report the age-associated emergence of unique epithelial cell states linked with thymic degeneration. These structural changes to the epithelial compartment could be linked to functional changes in the fibroblast compartment, specifically their upregulation of factors associated with inflammaging. These age-associated TEC populations perturbed thymic structure, bore transcriptional signatures of senescence and epithelial-to-mesenchymal transition and appear to act as a “sink” for epithelial growth factors. Their accumulation in the involuted thymus was exacerbated by acute injury and was associated with limited regenerative capacity compared to young mice. Thus, age-associated TECs represent a key feature of the involuted thymus that compromises organ function and may restrict regenerative capacity.

## RESULTS

### Emergence of atypical epithelial cell populations with thymic involution

Thymic involution is a well characterized chronic decline in thymic function over lifespan (*7, 9, 22, 23*). We found that total thymic cellularity declined in female mice across their lifespan (2, 6, 9, 12, 18 months) (**Fig. S1A**) coincident with morphological changes, such as a relative decrease in cortical-to-medullary ratio (**Fig. S1B-C**) (*24*). Quantification of the major structural cell lineages (TEC, EC and FB) by flow cytometry revealed little alteration in EC or FB, but a diminished TEC compartment that mirrors the overall loss of thymic cellularity, with a more severe loss in medullary TECs (mTECs) compared to cortical TECs (cTECs) despite the observed cortical thinning (**Fig. S1B-D**) (*25, 26*).

Recent reports have resolved the remarkable heterogeneity of TEC subsets (*16–20, 27*). To investigate the stromal changes in the thymic microenvironment associated with age-related thymic involution, we performed by single-cell sequencing of non-hematopoietic stromal cells (22,932 CD45^-^) from 2-month-old (2mo) or 18-month-old (18mo) female mice. Visualization of the data using dimensionality reduction methods revealed some broad stromal cell changes specific to the involuted thymus (**Fig. 1A, S2-3A**). To identify these cell types, the primary stromal cell lineages were defined based on transcription of lineage-specific genes: TECs with *Epcam*, *H2-Aa*; ECs with *Pecam1*, *Cdh5*; FBs with *Pdgfra*, alongside less abundant stromal cell types, including mesothelial cells (MECs) [*Upk3b*, *Nkain4*]; vascular smooth muscle cells (vSMCs) [*Acta2*, *Myl9*]; pericytes (PCs) [*Myl9*, *Acta2*] (**Figs. 1B, S3A, Data S1**); and extremely rare non-myelinating Schwann cells (nmSCs) [*Gfap*, *Ngfr* (p75) , *S100b*] (*28*). To more precisely define the heterogeneity of the major CD45^-^ structural compartments (epithelial, endothelial, fibroblast) we then subsampled and re-analyzed each major stromal cell population separately and developed ThymoSight (www.thymosight.org) to facilitate integration of previously published thymic sequencing data (*16, 17, 20, 21, 29*) with our own, which resulted in a dataset of 192,162 non-hematopoietic thymic stromal cells. Using this approach, we mapped the steady-state clusters within each cell lineage to publicly available datasets (**Fig. 1C-E, S3B-D**). Unsupervised clustering analysis distinguished the *Pdgfra*-expressing mesenchyme into 3 main groups, 2 of which were consistent with mouse capsular or human interlobular FBs (capsFB, based on expression of genes such as *Dpp4*, *Smpd3* and *Pi16*), and mouse medullary or human peri-lobular FBs (medFB, based on expression of markers such as *Ptn* and *Postn*) (*21, 30, 31*) (**Fig. 1D-E, S3B**). We also identified an intermediary subset of FB (intFB) marked by *Inmt3* and *Gpx3, t*hat did not map to the public gene signatures (**Fig. 1D-E, S3B**). Quantitative analysis of these populations indicated a small decrease in the frequencies of capsFB or intFB and an increase in medFB with age (**Fig. 1F**). We confirmed these changes by using the sequencing data to derive a flow cytometry panel, revealing an increase in the absolute number of medFBs (**Fig. 1G**). This age-related increase in fibroblasts may reflect that observed in other tissues where fibrosis is a key driver of functional loss (*32, 33*).

**Figure 1:**
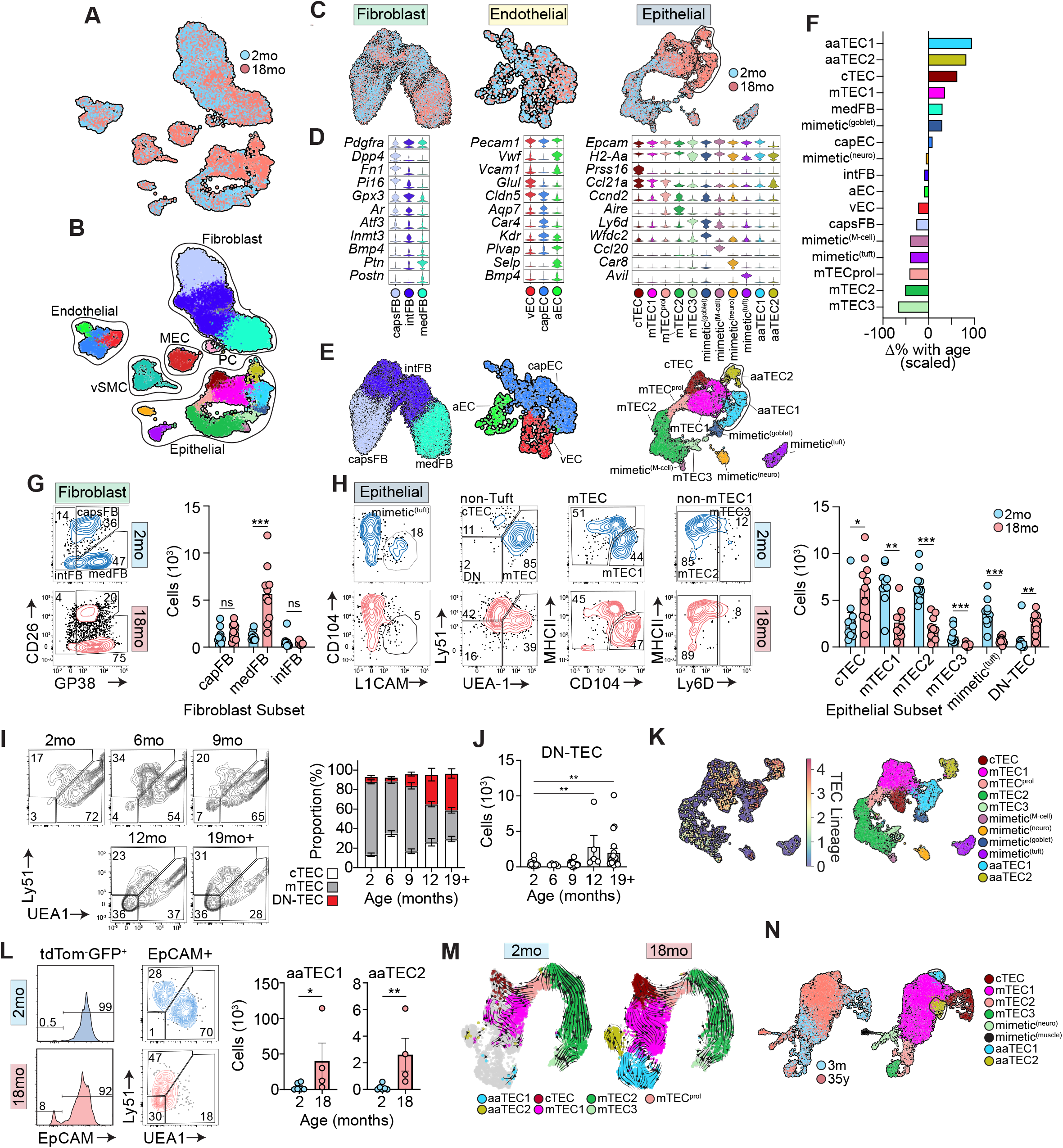
Emergence of atypical epithelial populations with age. **A-F**. Single cell RNA sequencing was performed on CD45^-^ cells isolated from the thymus of 2mo or 18mo female C57BL/6 mice. **A-B**, Uniform manifold approximation and projection (UMAP) of 22,932 CD45^-^ thymic cells annotated by age cohort (**A**), and cell type subset and outlined by cell type (**B**). **C-E**. **C**, UMAPs of individual structural compartments (endothelium, fibroblasts, epithelium) color-coded by age cohort. **D**, Violin plots highlighting key genes marking individual subsets within individual structural compartments. **E**, UMAPs of individual structural compartments color-coded by structural cell subset. n_EC_=1,661; n_FB_=13,240; n_TEC_=6,175. **F**, Scaled change in frequency for each individual structural cell subset with age. **G**, Concatenated flow cytometry plots and quantities for cell populations within the fibroblast lineage (n=10/age). **H**, Flow cytometry plots and quantities for cell populations within the epithelial lineage (first plot gated on CD45^-^EpCAM^+^) (n=10/age). **I**, Concatenated flow cytometry plots and graph highlighting the frequency of Ly51-UEA1- (DN-TEC) across lifespan. **J**, Number of DN-TEC across lifespan. **K**, Single cell RNA sequencing was performed on CD45^-^ cells isolated from 20mo *Foxn1^tdTom^*and age-matched wild type mice and integrated into the epithelial data described in **C-E**. UMAP of 8,505 cells of the epithelial compartment in the integrated data showing the overlaid expression of tdTomato (left). Isolated layer of the same UMAP showing only our TEC annotated subsets (right). **L**, Representative flow cytometry plots from *Foxn1^nTnG^* mice at the indicated ages, gated on tdTom^-^GFP^+^ cells, and number of GFP^+^tdTom^-^EpCAM^+^Ly51^-^UEA1^-^ aaTEC1 and GFP^+^tdTom^-^EpCAM^-^ aaTEC2 (n=4-6/age). **M**, RNA velocity on selected TEC populations in 2mo or 18mo mice. n_2mo_ = 1,989; n_18mo_ = 3,382. **N**, aaTEC1 and aaTEC2 gene signatures (top 30 marker genes from our mouse data converted to human orthologues, **Fig. S3, Data S3**) were overlaid on 3mo and 35yo human epithelial cells (n_TEC_ = 6,904) from single cell sequencing datasets generated and published in (*21*). Summary data represents mean ± SEM; *, p<0.05; **, p<0.01; ***, p<0.001 using the Mann-Whitney (G, H, L) or Kruskal-Wallis (J) tests.

Within the endothelium, three main clusters were identified (**Fig. 1D-E; S3C, Data S1**). Using gene signatures from an organ-wide murine EC atlas (*34*) these clusters were mapped to arterial (aEC), capillary (capEC) or venous EC (vEC) (**Fig. 1D-E; S3C, Data S1**). Less than 1% of all ECs showed enrichment for lymphatic markers (**Fig. S3C**), consistent with the scarcity of lymphatic ECs observed in sections or flow cytometric analysis of thymi from adult mice (*35*). Expression of *Plvap* and *Cldn5* delineated capEC (*36*), while aEC demonstrated the highest expression of *Vwf* and *Vcam1* (**Fig. 1D-E; S3C, Data S1**), genes that correlate with the vessel diameter. Venous ECs expressed high levels of P-selectin (*Selp*) (**Fig. 1D-E; S3C, Data S1**), indicative of thymic portal ECs (TPECs) (*37*), which have been linked to homing of hematopoietic progenitors in the thymus (*38–40*). Notably, *Bmp4*, which is produced by thymic ECs and is involved in regeneration after acute insult (*41*), was expressed highest by vECs (**Fig. 1D-E; S3C, Data S1**). We found little change in the frequencies of EC subsets or their number with age (**Fig. 1F, S4**).

Finally, TEC clusters were annotated based on the nomenclature and signatures derived from previously reported studies (*16–20, 42*) and mapped to nine subsets (**Fig. S3D, Data S1**): cTEC (based on, among others, expression of *Prss16, Psmb11,* and *Ly75*), mTEC1 (*Ccl21a, Itgb4,* and *Ly6a*), a proliferating mTEC subset (mTEC^prol^, *Ccnd2*), mTEC2 (*Aire*), and mTEC3 (*Ly6d* and *Spink5*). Notably, we found mimetic populations with neural (mimetic^neuro^, *Car8*), tuft-like (mimetic^tuft^; *Avil* and *L1cam*), goblet cell (mimetic^goblet^; *Spink5* and *Wfdc2*) and microfold (M)- like cells of the small intestine (mimetic^M-cell^; *Ccl20*); all of which correspond with the recently described transcriptional “mimetic” TECs (*27*) (**Fig. 1D-E; S3D, Data S1)**.

The reference data for each of these three structural cell lineages from the 2mo stroma were compared with the profiles from the 18mo involuted thymus, demonstrating considerable overlay (**Fig. 1C, Data S2**). In the case of ECs and FBs, no major new cell populations emerged with age, although there were transcriptional differences within existing cell populations, suggesting an age-associated change in cell state rather than cell type (**Fig. 1C, Data S2**). However, in the TEC compartment we observed two distinct age-associated epithelial cell types, referred henceforth as aaTEC1 and aaTEC2. These were apparent only in the 18mo (and not in the 2mo dataset) and could not be mapped with any published annotations (**Fig. 1C-E**). When we assessed the quantitative changes in cell subsets with age, we found an increased representation of cTEC and mTEC1 (consistent with previous reports (*20*)) but the two aaTEC populations exhibited the greatest changes with age (**Fig. 1F**). By flow cytometry we observed a marked increase of an atypical TEC population with age that expressed EpCAM but was negative for Ly51 and UEA1 (features of cTEC or mTEC, respectively) (**Fig. 1H**). This population of Ly51/UEA1 double negative (DN)TECs first emerged at 6-9 months and continued expanding across lifespan (**Fig. 1I-J**). This phenotype corresponded with the atypical aaTEC1 population which expressed few canonical cTEC or mTEC associated genes, while aaTEC2 also lacked expression of the canonical epithelial marker EpCAM (**Fig. 1D**). These data suggest that 2 unique epithelial cell types emerge in the involuting thymus with age that lack typical TEC features.

Given the paucity of expression of TEC genes by aaTEC, yet their clear association with other TEC populations (**Fig. 1B**), we sought to establish the lineage relationship between aaTECs and TECs. First, we created *Foxn1^Cre^*x R26-fl-Stop-fl tdTomato (*Foxn1^tdTom^*) reporter mice to drive constitutive tdTomato expression in all cells with a history of *Foxn1* expression, a master transcription factor in TEC (*43, 44*). Overlaying single cell RNA sequencing (scRNA-seq) data of 1,093 number of 20mo *Foxn1^tdTom^* TECs on the broader (7,412 WT cells: includes 6,175 18mo B6 and 1,237 20mo *Foxn1^WT^*controls) dataset revealed transcription of the tdTomato reporter gene across all lineages of TECs, including both aaTEC1 and aaTEC2 (**Fig. 1K**). The epithelial derivation of aaTEC was confirmed in a separate reporter strain, *Foxn1^Cre^* x *Rosa26^nTnG^* (*Foxn1^nTnG^*), where all cells express a nuclear-localized tdTomato reporter except those that have activated *Foxn1*, which instead express a nuclear-localized GFP protein (**Fig. S5**). Flow cytometric analysis of *Foxn1^nTnG^* mice confirmed the emergence of two atypical GFP^+^ TEC-derived populations with age: one population that was EpCAM^+^ but lacked canonical cTEC or mTEC markers, mirroring the DN-TEC/aaTEC1 population described above, and another that was EpCAM^-^, consistent with the aaTEC2 phenotype (**Fig. 1L**). These genetic approaches confirm a TEC origin of both aaTEC populations.

We next assessed their immediate precursors by performing unbiased RNA velocity analysis. Consistent with previous reports (*16, 20*), this approach demonstrated a clear lineage trajectory stemming from the mTEC^prol^ and mTEC1 populations and continuing into the more differentiated mTEC lineages (mTEC2 and mTEC3) in young mice. It is noData Sthat the recently described early and late TEC progenitor subsets overlaid with the mTEC1 cluster **(Fig. S3D**) (*29*). Upon thymic involution, mTEC1 also gave rise to aaTEC1, while aaTEC2 were derived from both aaTEC1 and mTEC1 (**Fig. 1M**). These data imply aaTEC represent end-stage epithelial cells arising and accumulating in the involuted thymus. Finally, to assess whether aaTEC are a feature of the human thymus, we assessed previously generated TEC scRNA-seq data from 3-month-old or 35-year-old samples (*21*). Using an aaTEC gene signature (**S3, Data S3**), we found populations corresponding to both aaTEC1 and aaTEC2 populations only in the adult dataset (**Fig. 1N, S6**). Overall, these data reveal the most prominent shifts in stromal cell type during thymic involution was the emergence of unique aaTEC populations.

### aaTEC form non-functional microenvironmental “scars” associated with partial EMT

The nuclear localization of the *Foxn1^nTnG^* reporter enabled cellular resolution of TEC by lightsheet imaging of whole cleared thymic lobes from young and old mice. As expected, we observed a relatively low density of GFP^+^ cells in the thymic cortex compared to the subcapsular and medullary regions (**Fig. 2A**). At the whole tissue level, the medulla formed a highly complex and interconnected structure in the young thymus (*45, 46*) that degenerated into isolated islets upon involution (**Fig. 2B, Supplemental Movie 1A-B**). Another striking feature of the involuted thymus, entirely absent in the young, was the emergence of zones of very high density GFP^+^ TEC (HD-TEC) clusters (**Fig. 2A-B**). These HD-TEC clusters formed band-like structures that were associated with the medulla of the involuted thymus. (**Fig. 2B, S7A, Supplemental Movie 1B**). Although the volume of cortex and medulla, and the number of cTECs and mTECs (calculated from whole-tissue imaging) declined with age, HD-TEC clusters emerged to compose a substantial volume and number of TEC (**Fig. 2C)**.

**Figure 2:**
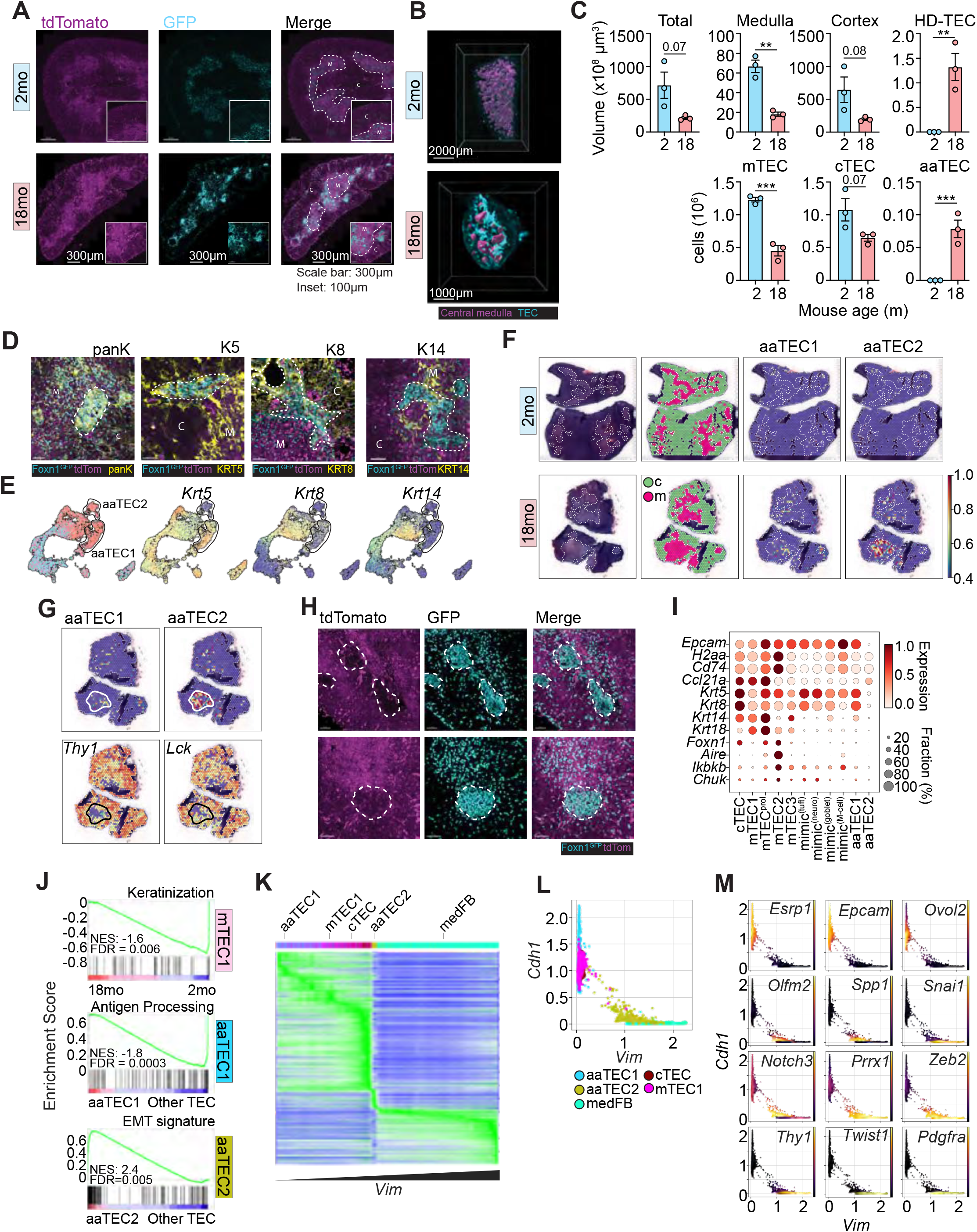
Emergence with age of distinct non-functional microenvironmental “scars” associated with EMT. **A**, Representative images of thymus in 2mo and 18mo *Foxn1^nTnG^*mice with medulla marked by dotted line. High density (HD-TEC) regions are apparent in aged but not young mice. **B,** Images of whole tissue light sheet imaging showing central medulla (magenta) in 2mo and 18mo female *Foxn1^nTnG^* mice with HD-TEC (cyan) only apparent in old mice. **C**, Quantification of total thymus volume, volume of cortex, medulla and HD-TEC regions and the number of mTEC, cTEC and HD-TEC calculated from whole tissue and confocal imaging. Summary data represents mean ± SEM; *, p<0.05; **, p<0.01; ***, p<0.001 using an unpaired t Test. **D**, Expression of pan-keratin and Keratin subunits 8, 5, and 14 on HD-TEC regions of thymus from aged mice. Scale bar: 50μm. **E**, Transcription of keratins subunits in the epithelial scRNA-seq dataset. n_TEC_ = 6,175. **F-G**, Visium spatial sequencing performed on 2mo or 18mo C57BL/6 thymus. **F,** Displayed are H&E sections, cortex and medulla identified by leiden clustering, and heatmaps of aaTEC1 and aaTEC2 signatures (top 10 differentially expressed genes for each subset vs other TEC, **Fig. S3, Data S3**) overlaid. **F,** Displayed are images of H&E stained sections, with cortical and medullary regions identified by leiden clustering, and heatmap of spatial transcriptome overlaid with signatures for aaTEC1 or aaTEC2. **G**, Expression of thymocyte markers *Thy1* and *Lck* overlaid on the 18mo spatial transcriptomics dataset. Outline represents thymocyte poor area overlaid onto heatmap showing aaTEC1 or aaTEC2 signatures. **H**, Two representative images showing td-Tomato and GFP expression with HD-TEC areas highlighted, with little or no td-Tomato^+^ cells. Scale bar: 50μm. **I**, Expression of key epithelial genes by various TEC subsets. **J**, GSEA pathway enrichment within mTEC1, aaTEC1, or aaTEC2 subsets (generated by comparing 18mo to 2mo mTEC1, **Data S4**; while for aaTEC1 and aaTEC2 generated by comparing to all other TECs, **Data S5**). **K**, Heatmap of 8,795 genes within cTEC, mTEC1, aaTEC1, aaTEC2, and medFB subsets ranked by vimentin expression. **L**, Scatterplot of *Cdh1* and *Vim* with cTEC, mTEC1, aaTEC1, aaTEC2, and medFB. **M**, Scatterplot of *Cdh1* and *Vim* transcription overlaid with expression of epithelial and mesenchymal genes.

Many GFP^+^ TEC in these high-density regions of the involuted thymus expressed keratin (**Fig. 2D, S7B**), confirming the retention of epithelial identity by these cells. To determine whether the HD-TEC structures were composed of aaTEC, we surveyed the expression of keratin subunits distinguishing aaTEC from other TEC populations (**Fig. 2D-E**). We found that most GFP^+^ TEC in these regions were K5^+^, some were K8^+^ and none were K14^+^ (**Fig. 2D-E**); a profile corresponding to transcriptional profile of aaTEC1 (**Fig. 2E**). Spatial transcriptomics comparing thymi from 2mo to 18mo mice confirmed aaTEC1 and aaTEC2 signatures formed clusters that were only detected in the involuted thymus (**Fig. 2F**). The medullary distribution (as assessed by H&E staining) of these signatures resembled that of HD-TEC clusters (**Fig. 2F, S3, Data S3**). However, there was a marked lack of thymocyte transcripts in the aaTEC zones (**Fig. 2G**). Consistent with this observation, HD-TEC clusters in the involuted thymus of *Foxn1^nTnG^*mice excluded other tdTomato^+^ cells (**Fig. 2H, S7**), which given its ubiquitous expression in non- *Foxn1* derived cells, will largely mark thymocytes. These data suggest that aaTEC microenvironments were thymocyte “deserts” that do not support thymocyte differentiation and are deprived of thymic crosstalk factors. Consistent with this notion, we found that aaTEC express low levels of *Foxn1*, *Aire* or downstream NFκB target genes (**Fig. 2I, S7C-D**). Moreover, in accord with the scRNAseq analyses, imaging confirmed that aaTEC did not represent clusters of mimetic cells, lacking expression of markers of tuft cells or M-cells (**Fig. S7E**). Taken together, these data strongly suggest functional equivalence between DN-TEC identified through flow cytometry, HD-TEC identified by imaging approaches, and aaTEC identified through scRNA-seq. Although not fibrotic themselves, these aberrant high-density aaTEC “scars” mirror mesenchymal scarring found in other tissue with age and may be responsible for diminishing overall thymic function (*32, 33, 47*).

The unusual morphology and microenvironment formed by aaTEC prompted us to assess the functional changes in TECs with age. Gene set enrichment analysis (GSEA) (*48, 49*) revealed loss of the keratinization pathway in mTEC1 with age, and loss of antigen presentation within aaTEC1 (vs all other TEC) (**Fig. 2J, Data S4-5**), consistent with the progressive loss of epithelial function upon differentiation into aaTEC. Furthermore, one of the most highly enriched pathways in aaTEC2 (vs all other TEC) was the hallmark EMT gene signature (**Fig. 2J, Data S5**), highlighting a loss of epithelial and gain of mesenchymal identity. A heatmap of gene expression across 8,795 EMT-related genes in ascending order according to expression of vimentin, as previously described (*50*), further suggests that aaTEC2 lies in a liminal zone between TECs and fibroblasts (**Fig. 2K**). Scatterplots based on E-cadherin (*Cdh1*) and vimentin (*Vim*) transcription (as prototypical epithelial and mesenchymal markers, respectively) overlaid with expression of archetypal epithelial or mesenchymal genes further suggested that aaTEC2 lost their epithelial state and partially gained mesenchymal traits (**Fig. 2L-M**). These observations are most consistent with a partial EMT (pEMT) population that parallels a senescence-associated pEMT tubular epithelial population identified in kidney fibrosis (*51*), although we do not exclude the possibility that some aaTEC2 undergo a full EMT (*52*). Given the intrinsic link between EMT, fibrosis, and senescence (*32, 53, 54*), these data suggest that aaTEC form unique, non-functional microenvironments in the involuted thymus associated with senescence and EMT.

### aaTEC expansion and delayed thymic regeneration following acute injury in old mice

The thymus is extremely sensitive to injury but has substantial capacity for repair. The ability of the thymus to regenerate is thought to decline with age yet the mechanisms of this deficit are poorly understood (*7–9*). We found that 1-2mo mice subjected to sub-lethal total body irradiation (TBI) had fully recovered thymic cellularity by day 28 after TBI (**Fig. 3A**). By contrast, aged mice exhibited a significant delay in the restoration of thymic cellularity and did not approach normal levels until approximately day 42 (**Fig. 3A**). Plotting thymic size after damage relative to baseline cellularity showed that restoration of thymic cellularity during the regenerative phase was impaired in aged mice (**Fig. 3B**). Histomorphological analysis confirmed these findings with similar depletion of thymocytes, accumulation of cellular debris, and granulation tissue formation 1 day after injury between aged cohorts, but evidence of fibrosis, dystrophic calcification and occasional dyskeratotic epithelial cells only in 18mo mice, suggesting a relative dysfunction in the regenerative process with age (**Fig. S8A**).

**Figure 3.**
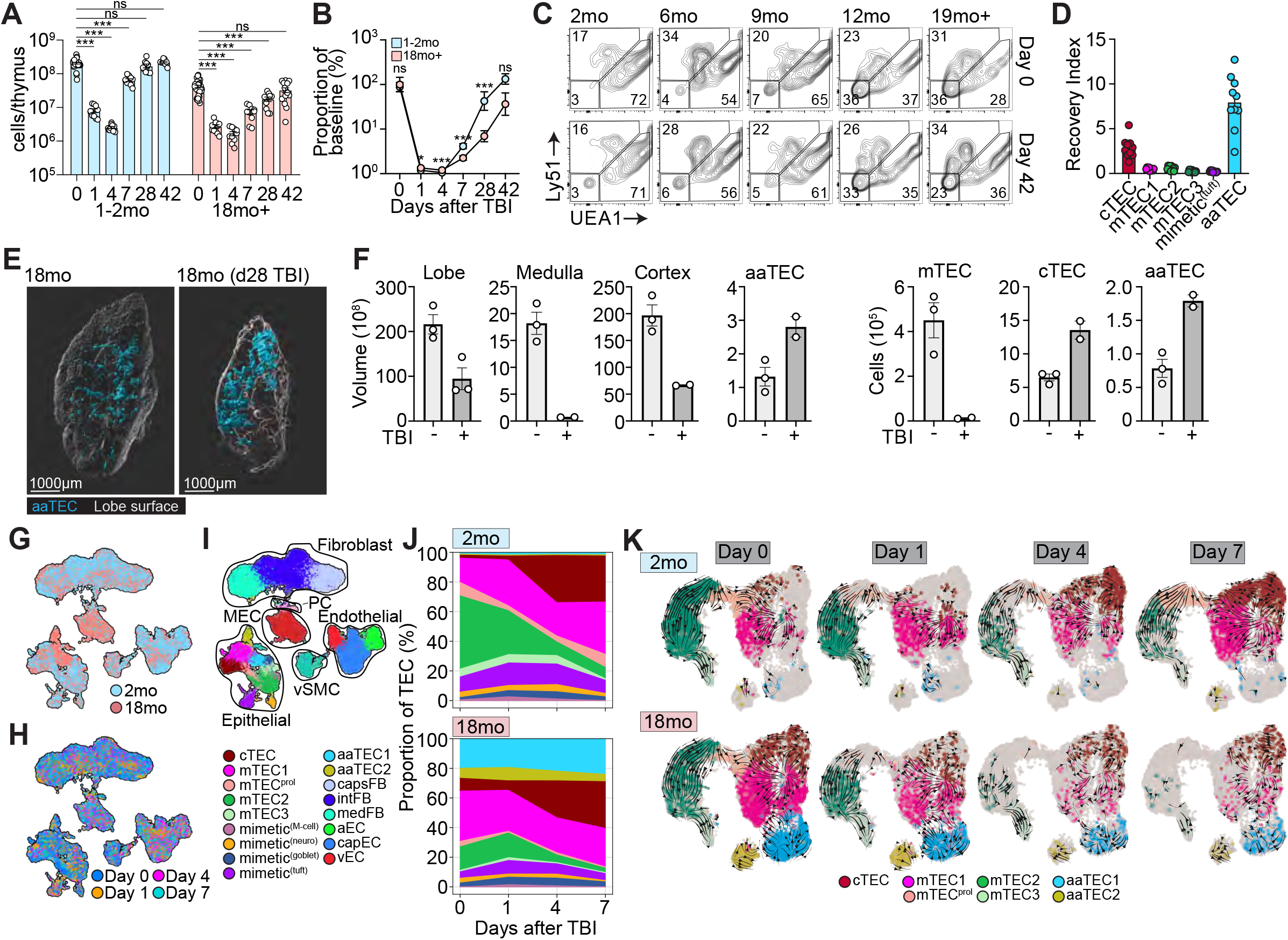
Aging negatively impacts thymic regeneration. **A-B,** 2mo or 18mo female C57BL/6 mice were given a sublethal dose of total body irradiation (550cGy) and the thymus was assessed at the indicated timepoints. **A,** Total thymic cellularity. **B,** Proportion of total thymic cellularity at the indicated timepoint as a function of steady-state age-matched cellularity. **C,** Flow cytometry plots (gated on CD45^-^EpCAM^+^) showing aaTEC1 at day 42 after TBI in C57BL/6 mice at the indicated ages. **D**, Depletion and recovery of indicated populations were quantified by flow cytometry over the first 7 days after TBI in 2mo or 18mo mice and area under the curve was calculated. An aging index was generated by calculating the ratio of aged to young AUC for each indicated population (n=10/celltype). **E**, Whole tissue imaging of 12mo Foxn1^nTnG^ mice at baseline or 28 days after TBI. **F**, Total volume and volume of cortex, medulla and aaTEC regions, as well as the number of cTECs, mTECs and aaTECs of thymic right lobe. **G-J**, Single cell RNA sequencing was performed on CD45^-^ cells isolated from 2mo or 18mo thymus at baseline (day 0) and days 1, 4 and 7 after TBI. UMAP of 81,241 CD45^-^ cells annotated by age cohort (**G**), day after TBI (**H**), or structural cell subset mapped from **Fig. 1B-S9** (**I**). **J** Associated frequency analysis of all TEC subsets after TBI within each age cohort. **K**, RNA velocity analysis on selected TEC subsets at days 0, 1, 4, and 7 after TBI in 2mo or 18mo mice. Summary data represents mean ± SEM; *, p<0.05; **, p<0.01; ***, p<0.001 using the Kruskal-Wallis (A, D), Mann-Whitney (B), or unpaired t (F) tests.

To assess the stromal compartments that orchestrate thymic regeneration, we performed flow cytometric analysis of endothelial, mesenchymal, and epithelial stromal cell populations before and after regeneration. Although there were few major differences in the early response of young versus aged mice in most populations (**Fig. S8B**), one noData Sexception was the persistence of the Ly51^-^UEA1^-^ DN aaTEC1, which remained a prominent feature of the regenerated thymus of old mice (**Fig. 3C-D, S8B)**. Whole organ imaging analysis of aged *Foxn1^nTnG^*mice 28 days after TBI revealed prominent high-density aaTEC regions after damage (**Fig. 3E, Supplemental Movie 2**). Quantification revealed that, despite the thymus remaining smaller than pre-damage, aaTEC composed an increased volume and number, , indicating a substantial preferential increase in aaTEC coincident with the impaired thymic regeneration of aged mice (**Fig. 3E-F**).

To comprehensively explore the mechanisms of the damage response and impaired regeneration in aged thymic stroma, we analyzed 58,309 CD45^-^ non-hematopoietic cells from the thymus of 2mo and 18mo by scRNA-seq at days 0,1, 4 and 7 after TBI and annotated them based on baseline (day 0) subset signatures, prior to integrating them together (**Fig. 3G-K, S2, S9 Data S1-3**). Age-associated TEC populations remained a major feature of the aged thymic recovery after damage **(Fig. 3J**). RNA velocity analysis implied that, in 2mo mice, the reemergence of differentiated mTEC populations stemmed largely from the mTEC1 and mTEC^prol^ populations (**Fig. 3K**). In contrast, in 18mo mice there was a skewing in the inferred direction from the mTEC1/mTEC^prol^ population towards the aaTEC1 and aaTEC2 rather than the mTEC2/3 differentiated mTEC subsets (**Fig. 3K**), consistent with findings by others (*20*). Therefore, the emergence of aaTEC perturbs normal TEC differentiation in the early stages of regeneration.

### Inflammaging drives the emergence of aaTEC, which co-opt tonic signals from TECs at baseline and reparative cues after damage

We next sought to understand the molecular changes that may underlie the functional changes to the thymus with age. We first used GSEA to identify pathways associated with age-dependent transcriptional responses within each specific structural cell subset and coupled it with network enrichment analysis using Cytoscape (*55*) to integrate the GSEA results into networks sharing common gene sets. Pathways with significant enrichment across at least one population with age were broadly grouped into 8 biological categories: antigen processing and presentation; immune function; mitochondrial function; metabolism; proteostasis; extracellular matrix interactions; solute sensing; and cell differentiation and function (**Figs. 4A, S10, Data S4**). Notably, many of these categories corresponded with the main hallmarks of aging that have been identified across organisms and tissues (*56–58*). We found a broad decrease in the transcription of genes within pathways associated with mitochondrial function and metabolism across all structural cell lineages including fibroblast, endothelial and epithelial subsets (**Fig. 4A, S9, Data S4**), consistent with a link between mitochondrial function, aging and senescence (*59, 60*). In fibroblasts, evidence of reduced mitochondrial function was attended by upregulation of genes associated with immune function (including antigen processing and presentation), consistent with the role of these cells and pathways in the induction of inflammaging or the senescence-associated secretory phenotype (SASP) (*61–64*). By contrast, a decrease in pathways associated with proteostasis was largely restricted to epithelial cell populations (**Fig. 4A, S9, Data S4**).

**Figure 4:**
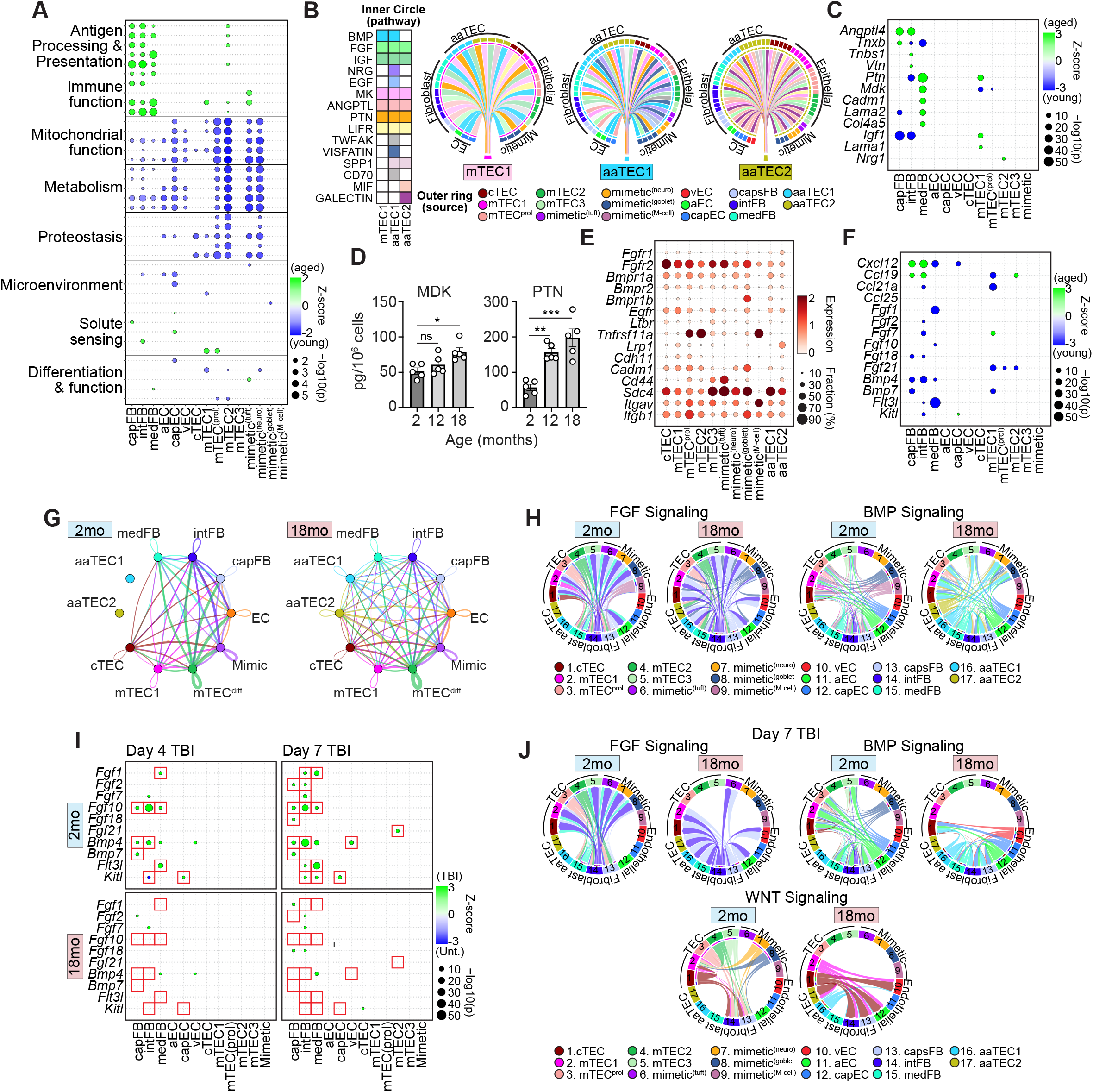
Muted transcriptional response to irradiation in the aged thymic stroma linked to aaTEC disruption of trophic signals. **A,** GSEA pathway analysis was performed for each subset based on differentially expressed genes in 2mo versus 18mo mice (**Data S4**). Dotplot of top 5 pathways within each category. **B,** CellChat interaction analysis between stromal cell populations with mTEC1, aaTEC1 or aaTEC2 as cellular receivers. Outer rings represent cell source and correspond to the population coloring used throughout the manuscript. Inner circle represent significantly enriched pathways targeting each individual cell population. Matrix on the left represents all significantly enriched pathways targeting either mTEC1, aaTEC1 or aaTEC2 and colored according to which cell the pathway interacts (colored according to pathway). **C**, Differential expression of inflammaging/SASP genes between 2mo and 18mo mice by the indicated stromal cell subsets. **D**, Amounts of PTN and MK in thymus at 2mo, 12mo or 18mo female C57BL/6 mice (n=5/age). Summary data represents mean ± SEM, each dot represents an individual biological replicate; *, p<0.05; **, p<0.01; ***, p<0.001 using the Kruskal-Wallis Test. **E**, Differential expression of key thymopoietic and epithelial growth factors between 2mo or 18mo mice across stromal cell subsets. **F**, Expression of receptors for epithelial growth factors and EMT-driving factors across stromal cell subsets. **G,** Circos plot summarizing number of interactions (derived from CellChat analysis) among stromal cell lineages. **H,** Chord Diagram interaction analysis of FGF and BMP signaling pathways in 2mo or 18mo mice at baseline. **I**, Differential expression across stromal cell subsets of key thymopoietic and epithelial growth factors between 2mo or 18mo female C57BL/6 mice at days 4 and 7 after TBI. **J,** Chord Diagram interaction analysis of FGF, BMP and WNT signaling pathways in 2mo or 18mo mice at day 7 after TBI.

Since the approach above involved direct comparison of cell populations from young and aged mice, aaTECs could not be assayed. Therefore, we next interrogated intercellular communication networks using CellChat (*65*) to infer cell-cell interactions. Significant enrichment of pathways (colored according to the matrix, left panel) is shown by the vectors connecting all stromal “source” populations with the “receiver” populations, focusing on aaTEC1, aaTEC2 or mTEC1 (as a likely cellular source of aaTECs) (**Fig 4B**). We found significant enrichment of several growth factor pathways, including those known to drive TEC growth (BMP, FGF, EGF, WNT), as well as pathways implicated in promoting EMT (ANGPTL, MDK, PTN, GALECTIN) (*66–71*) (**Fig. 4B**). Next, we looked at differential expression with age in genes selected by comparing GSEA EMT signature genes (**Fig. 2J**) with those represented within the upregulated inflammaging signatures in fibroblasts identified in **Fig. 4A.** We reasoned that these genes and pathways may be key drivers of atypical aaTEC differentiation. This manual gene curation revealed a broad upregulation of hallmark EMT inducing genes, especially amongst fibroblasts, including the inflammaging/SASP-associated factors *Angptl4, Ptn, Mdk*, and *Lgals1* (**Fig. 4C**), potentially explaining the emergence of aaTEC themselves (*66–71*). Consistent with this hypothesis, the amounts of PTN and MDK increased in the thymus with age (**Fig. 4D**) and, along with many other TECs, aaTECs expressed abundant EMT-related integrins and *Sdc4*, which can act as a receptor for multiple EMT ligands, including PTN, MK and ANGPTL4 (*72*) (**Fig. 4E**).

Concurrent with the upregulation of inflammaging/SASP and EMT programs, we also found decreased transcription of thymopoietic factors (e.g. *Flt3l* and *Kitl*), as well as members of the FGF and BMP epithelial growth factor pathways (**Fig. 4F**). Age-associated TEC1 and aaTEC2 expressed receptors for BMP, FGFs and EGF pathways, but not those for lymphotoxin or RANKL (*Ltbr* and *Tnfrsf11a*, respectively), canonical drivers of mTEC differentiation (*73*) (**Fig. 4E**). Interaction maps among cells based on expression of receptor-ligand pairs demonstrated a clear skewing of interactions with age toward aaTECs (**Fig. 4G**). This co-opting of trophic TEC signals, which include the FGF and BMP pathways, suggests that aaTECs draw these pro-growth ifactors away from conventional TEC with age (**Fig. 4H**). We therefore found both reduced production of thymopoietic growth factors as well as an increase in their signaling through aaTECs at the expense of conventional TECs (**Fig. 4F-H**).

In young mice, we found a broad upregulation of endogenous epithelial regenerative factors after damage such as *Fgf7, Fgf10, Fgf21, and Bmp4*, as well as thymopoietic factors like *Flt3l* and Kitl (**Fig. 4I**). In contrast, aged mice do not activate these same regenerative programs (**Fig. 4I**). Moreover, in addition to reduced expression of these regenerative pathways, our interaction analysis suggested BMP, FGF, and WNT interactions are skewed toward aaTECs in the aged, damaged thymus (**Fig. 4J**) (*7–9*).

## DISCUSSION

Here, we define age-associated changes to the microenvironment in the involuting thymus that impairs function in two ways. Firstly, atypical aaTECs form high density epithelial clusters, devoid of thymocytes. Therefore, the accretion of aaTEC regions directly contributes to the loss of functional thymic tissue with age and, given that these regions expand after damage, exacerbates injury-induced atrophy. Secondly, we also found evidence that the emergence of aaTEC perturbs the network of growth factors supporting stromal cell function and thymocyte differentiation, likely constituting an additional impediment to thymic function.

Despite the relatively early emergence of aaTECs, their genesis appears linked to hallmarks of aging. Both aaTEC populations accrued with age, starting at 6 months, and could be broadly divided into EpCAM^+^ and EpCAM^-^ populations. Genetic approaches demonstrated that these aaTEC populations were derived from a *Foxn1*^+^ precursor, yet both had lost expression of canonical markers of cTECs and mTECs. RNA velocity analysis suggests that both aaTEC1 and aaTEC2 derive from mTEC1, consistent with the progenitor-like phonotype of the latter (*16, 20*); however, lineage-tracing approaches will provide a more stringent test of this hypothesis. The molecular drivers of aaTEC likely involve the main hallmarks of aging (*56, 57*), reflected by a loss in mitochondrial, metabolic, and proteostasis programs in TECs, and increase in pathways associated with inflammaging or senescence-associated secretory phenotype (SASP) in fibroblasts (*57*). This latter finding is particularly noData Sgiven the likely role of inflammaging and SASP in driving EMT in aged tissues (*53, 54*). Our findings are especially noData Sgiven that one of the main features of the aging thymus is the replacement of functional tissue with fat (*5, 74*), with some evidence that this may be triggered by EMT (*75*), and adipogenesis itself can directly drive loss of thymic function (*76*). These data also support recent observations that age-associated changes are organ-specific and cell lineage specific (e.g. senescence-associated inflammaging impacting on muscle regeneration) (*32, 57, 58*). Our findings are consistent with the notion that aaTEC represent a thymus-specific manifestation of these programs.

These observations suggest that the defective response and recovery to acute damage with age could be due to: 1) failure of aged subsets to regulate necessary genes in the regenerative phase; 2) broad upregulation of an EMT-driving program leading to rapid re-emergence of aaTEC; and 3) those regenerative responses that are mounted are diverted to sustaining aaTEC rather than typical differentiated mTEC subsets. Notably, this seems to be at least partially driven by functional changes within the fibroblast compartment, and in particular their upregulation of genes associated with inflammaging/SASP. In summary, these studies offer insight into responses of specific structural cell subsets to aging and acute damage. Furthermore, the discovery of aaTEC along with the functional changes in fibroblasts with age consistent with inflammaging/SASP provide a therapeutic target for improving T cell immunity more broadly. Age-associated TECs therefore constitute a nexus of stromal cell dysfunction in thymic involution and impaired regeneration.

## ACKNOWLEDGEMENTS

We thank Dr. Katia Manova-Todorova, Eric Chan, Radoslaw Junka, Ning Fan, and Afsar Barlas from the Molecular Cytology Core Facility at Memorial Sloan-Kettering Cancer Center for their support and expert advice in the preparation and analysis of immunohistochemistry data. We also thank the Flow Cytometry and Comparative Medicine Core Facilities, as well as support of the Immunotherapy Integrated Research Center at the Fred Hutchinson Cancer Center. We are also grateful for the support of the WEHI Centre for Dynamic Imaging, Flow Cytometry facility and Bioservices facility, in particular L. Whitehead, P. Rajasekhar, T. Boudier S., Monard, Z. Arnold, H. Marks, T. Ballinger and S. Holloway.

## FUNDING SOURCES

This research was supported by National Institutes of Health award numbers R01-CA228358 (M.R.M.vdB), R01-CA228308 (M.R.M.vdB), R01-HL123340 (M.R.M.vdB), R01-HL147584 (M.R.M.vdB), P01-CA023766 (M.R.M.vdB), P01-AG052359 (J.A.D and M.R.M.vdB), R01-HL145276 (J.A.D.), R01-HL165673 (J.A.D.), U01-AI70035 (J.A.D.), R00-CA176376 (J.A.D.), and the NCI Cancer Center Support Grants P30-CA015704 (Fred Hutchinson Cancer Center) and P30 CA008748 (Memorial Sloan Kettering Cancer Center). D.H.D.G. was funded by Australian NHMRC Fellowships 1090236 and 1158024; NHMRC Project/Ideas Grants 1078763, 1121325, and 1187367 and Cancer Council of Victoria Grants-in-Aid 1146518 and 1102104 and through the Victorian State Government Operational Infrastructure Support and Australian Government NHMRC IRIISS. M.R.M.vdB also received support from the Starr Cancer Consortium, the Tri-Institutional Stem Cell Initiative, The Lymphoma Foundation, The Susan and Peter Solomon Divisional Genomics Program, Cycle for Survival, and the Parker Institute for Cancer Immunotherapy. J.A.D received additional support from a Scholar Award from the American Society of Hematology (J.A.D.); the Mechtild Harf (John Hansen) Award from the DKMS Foundation for Giving Life (J.A.D.); the Cuyamaca Foundation (J.A.D.); and the Bezos Family Foundation (J.A.D.). LJ received support from the European Molecular Biology Organization (ALTF-431-2017) and the MSK Sawiris Foundation. K.Z. is supported by a University of Melbourne Research Training Program Scholarship.

## AUTHOR CONTRIBUTIONS

AIK, LJ, KZ, EV, DHDG, JAD, MRMvdB conceived and designed the study. NRM, KLR, DHDG, JAD, MRMvdB acquired funding. LJ, AIK planned research activities. AIK, LS, LJ, LM curated data. AIK, LS, LJ performed initial analysis. AIK, LS, LJ, MS, RS, ALCG provided code and performed computational and statistical analysis of single-cell and bulk RNA-seq data. AIK created the R shiny interactive app and KC assisted with website development. LJ, KZ, JT, EV, KVA, FM analyzed flow cytometric data. AEF, NRM provided scRNAseq on *Foxn1^tdTom^.* KVA analyzed and interpreted H&E sections. LJ, KZ, AEF, DG, KC, JMS, EV, KVA, JT, AL, KN, NL, RG, FM, HA, SY, MBdS, MD, VCW, KLR, SDW, and BG planned and performed experiments. DP, VCW, NRM, DHDG, JAD, MRMvdB supervised study. LJ, AIK, KZ, SY, DHDG, JAD prepared figures and visualized data. LJ, AIK, KZ, DHDG, JAD, MRMvdB wrote the initial draft. All authors reviewed and approved the manuscript.

## DECLARATION OF INTERESTS

Dr. Marcel van den Brink has received research support and stock options from Seres Therapeutics and stock options from Notch Therapeutics and Pluto Therapeutics; he has received royalties from Wolters Kluwer; has consulted, received honorarium from or participated in advisory boards for Seres Therapeutics, WindMIL Therapeutics, Rheos Medicines, Merck & Co, Inc., Magenta Therapeutics, Frazier Healthcare Partners, Nektar Therapeutics, Notch Therapeutics, Forty Seven Inc., Priothera, Ceramedix, Lygenesis, Pluto Therapeutics, GlaskoSmithKline, Da Volterra, Vor BioPharma, Novartis (Spouse), Synthekine (Spouse), and Beigene (Spouse); he has IP Licensing with Seres Therapeutics and Juno Therapeutics; and holds a fiduciary role on the Foundation Board of DKMS (a nonprofit organization). The Walter and Eliza Hall Institute of Medical Research receives milestone and royalty payments related to venetoclax. Employees are entitled to receive benefits related to these payments; D.H.D.G reports receiving benefits. DHDG has received research funding from Servier.

## Supplementary Figures

**Fig. S1:**
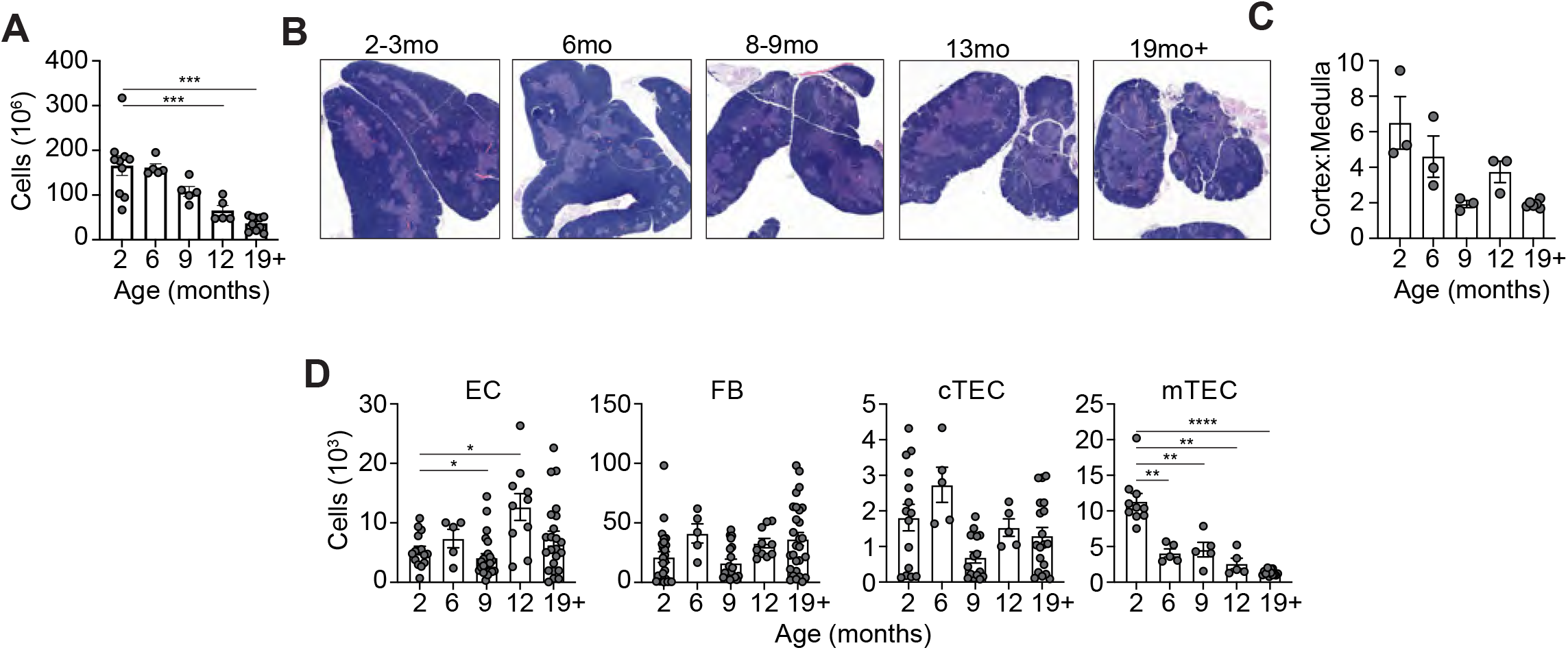
**A**, Thymic cellularity of female C57BL/6 mice at 2, 6, 9, 12 or 19+ months old. **B-C,** Representative images of hematoxylin and eosin (H&E) stained mouse thymi from 2, 6, 8-9, 13, and 19+mo female C57BL/6 mice (**B**) used to calculate ratio of cortical (dark) to medullary (light) region (**C**). In (**C**), each dot represents a biological replicate. **D,** Flow cytometric analysis of enzymatically-digested thymus and absolute cell numbers for major cell types (TECs; cTECs and mTECs; ECs and FBs) in 2, 6, 9, 12, and 19+-mo female C57BL/6 mice. Summary data represents mean ± SEM; *, p<0.05; **, p<0.01; ***, p<0.001 using a one way ANOVA.

**Fig. S2:**
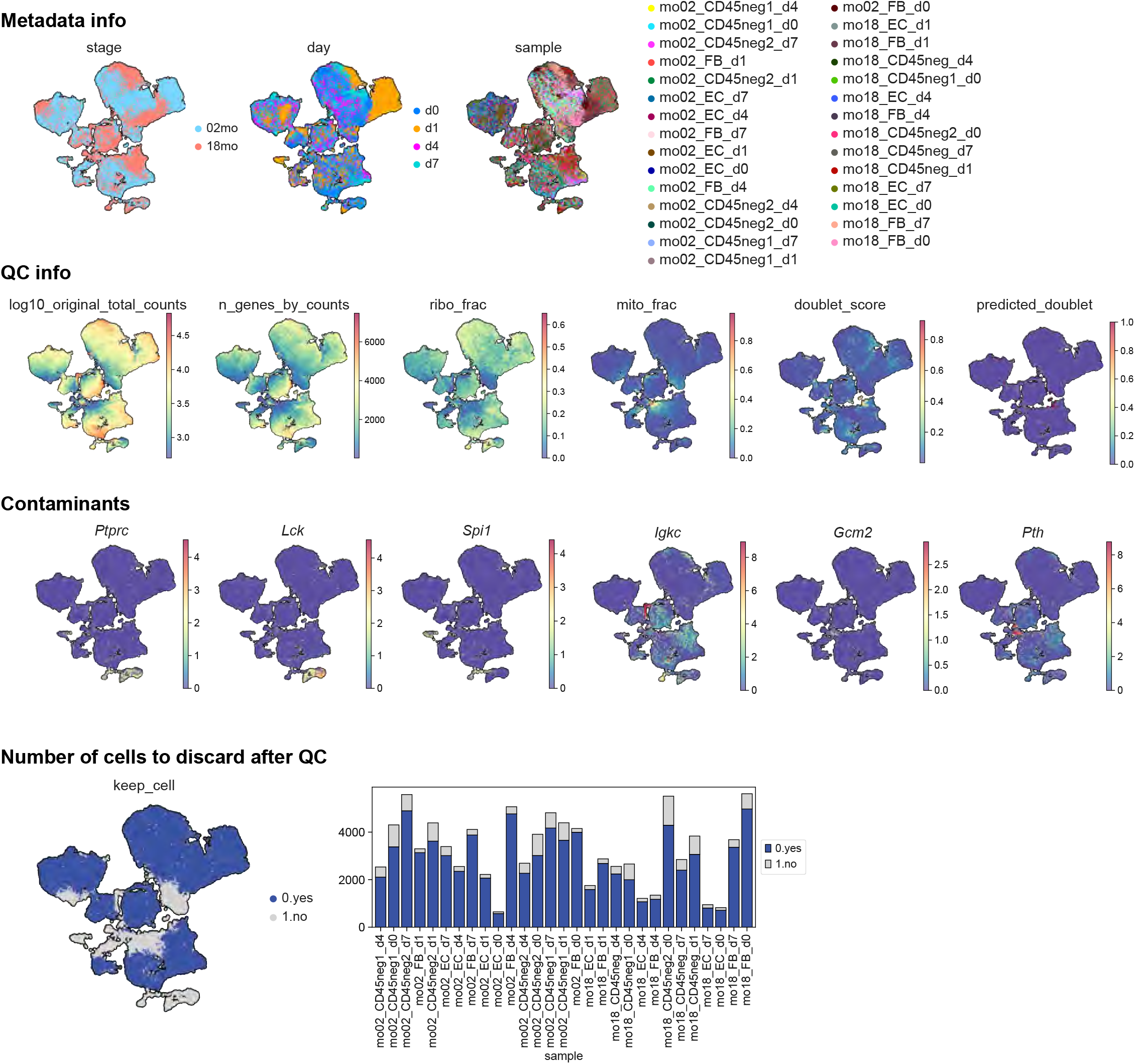
**A**, UMAPs of 93,738 CD45^-^ cells from 2mo or 18mo thymus at baseline (day 0) and at days 1, 4 and 7 after TBI prior to quality control. UMAPs in sequence display age cohort, day after TBI and origin sample (Metadata info); total counts in *log*_10_ scale, number of genes, ribosomal fraction, and mitochondrial fraction per cell (QC info); CD45^+^ and parathyroid contaminating cells expressing *Ptprc*, *Lck*, *Spi1*, *Igkc and Gcm2*, *Pth* genes respectively (Contaminants). **B**, UMAP and associated stacked barplot representation of filtered out cells on a per sample basis. Cells retained for further processing are shown in blue, while cells removed are shown in grey.

**Fig. S3:**
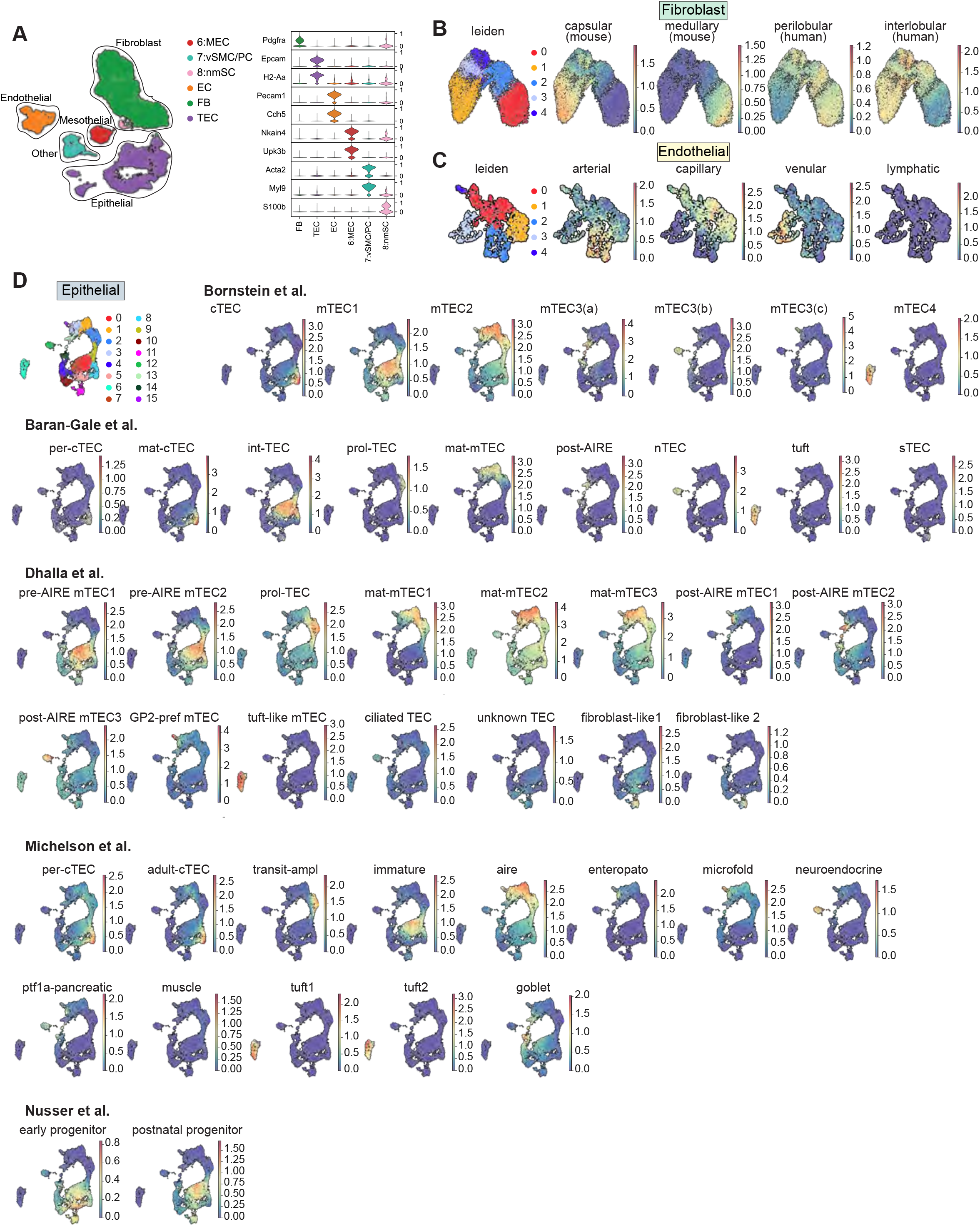
**A**, Broad structural cell subsets were annotated based on expression of canonical markers such as *Pdgfra, Epcam, H2-aa, Pecam*, and *Cdh5.* **B**, Leiden clustering of fibroblast subsets and signatures for murine capsular and medullary fibroblasts FBs and human perilobular and interlobular FBs based on previously published datasets (*21, 30, 31*). **C**, Leiden clustering of endothelial subsets and signatures for arterial, capillary, venular, and lymphatic ECs based on previously published datasets (*34*). **D**, Thymic epithelial cell signatures were compared to previously published literature and overlaid on our sequencing dataset (*16, 17, 20, 27*).

**Fig. S4:**
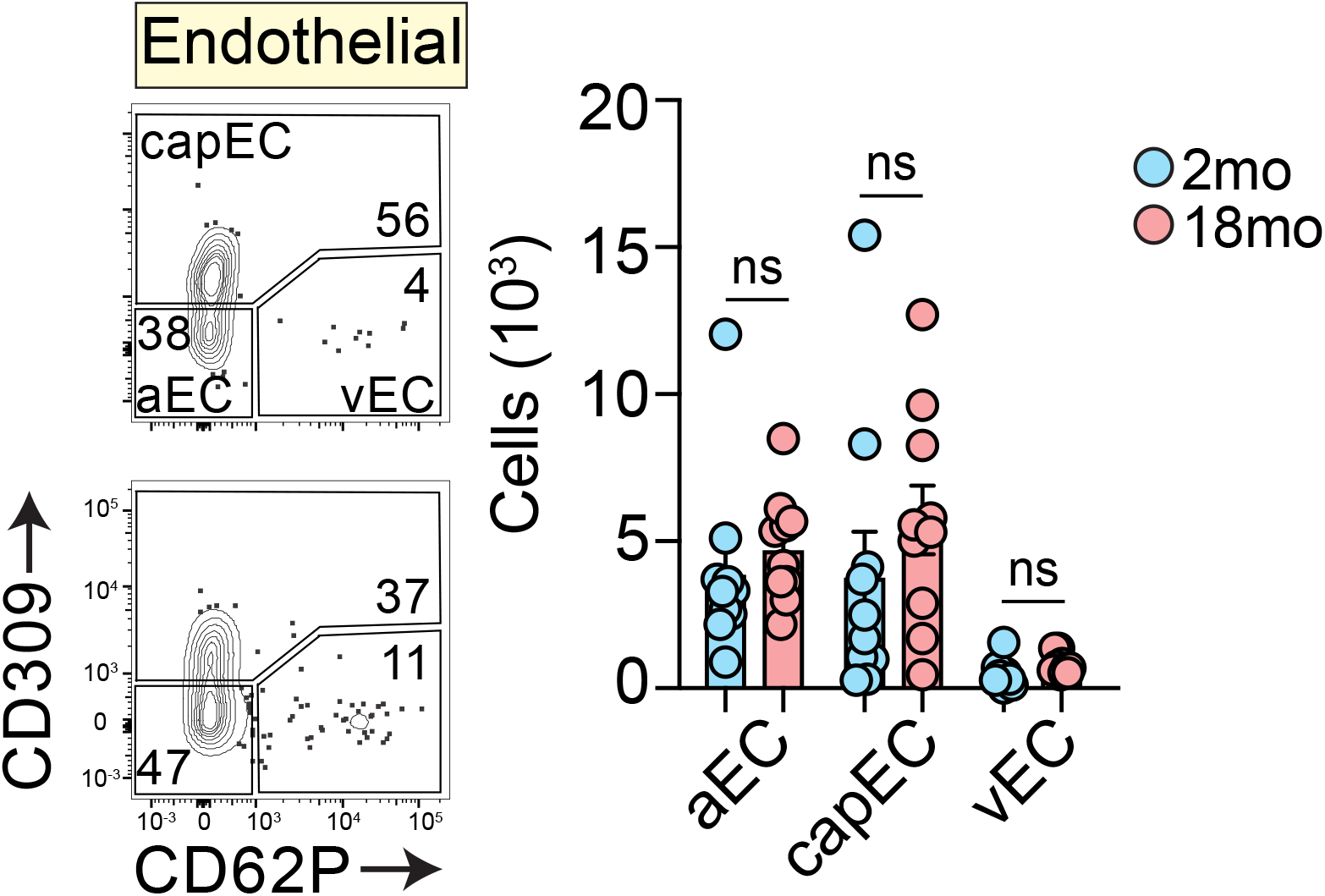
Concatenated flow cytometry plots and quantities for cell populations within the endothelial cell lineage (n=10/age). Data represents mean ± SEM; *, p<0.05; **, p<0.01; ***, p<0.001 using a Mann-Whitney test.

**Fig. S5:**
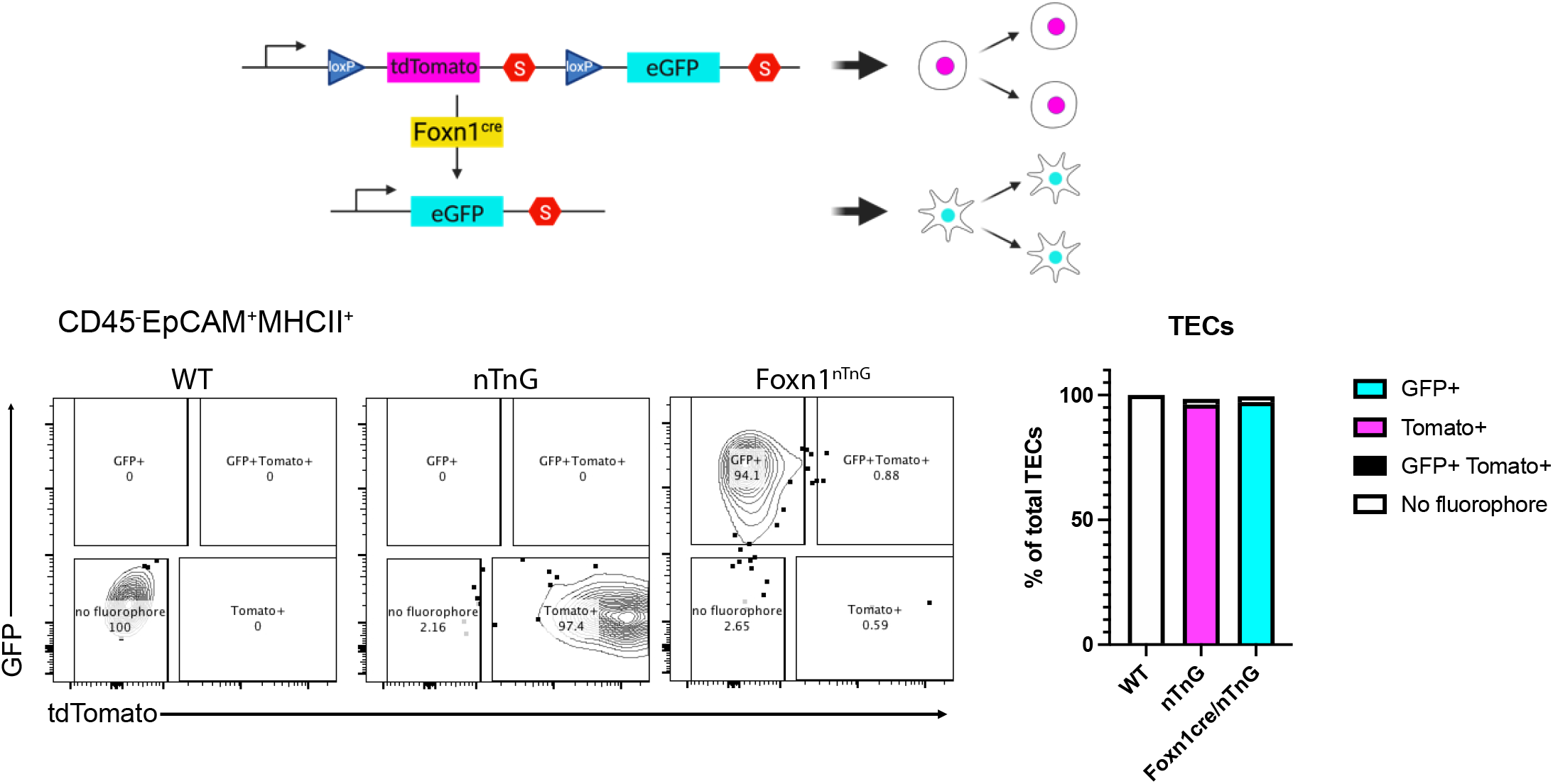
Generation of *Foxn1^nTnG^ ^mice^*. ROSA^nT-nG^ (nT/nG) mice (PMID: 24022199) were intercrossed with *Foxn1^Cre^* mice (PMID: 19234195). Representative flow cytometric plots of TEC from 11 weeks old WT, nT/nG and *Foxn1^nTnG^* show specific detection of GFP in nearly all TEC only in the latter strain. Quantification of the relative proportions of TEC expressing the reporters are shown in the bar graph on the right (n= 2 to 3 from 2 experiments).

**Fig. S6:**
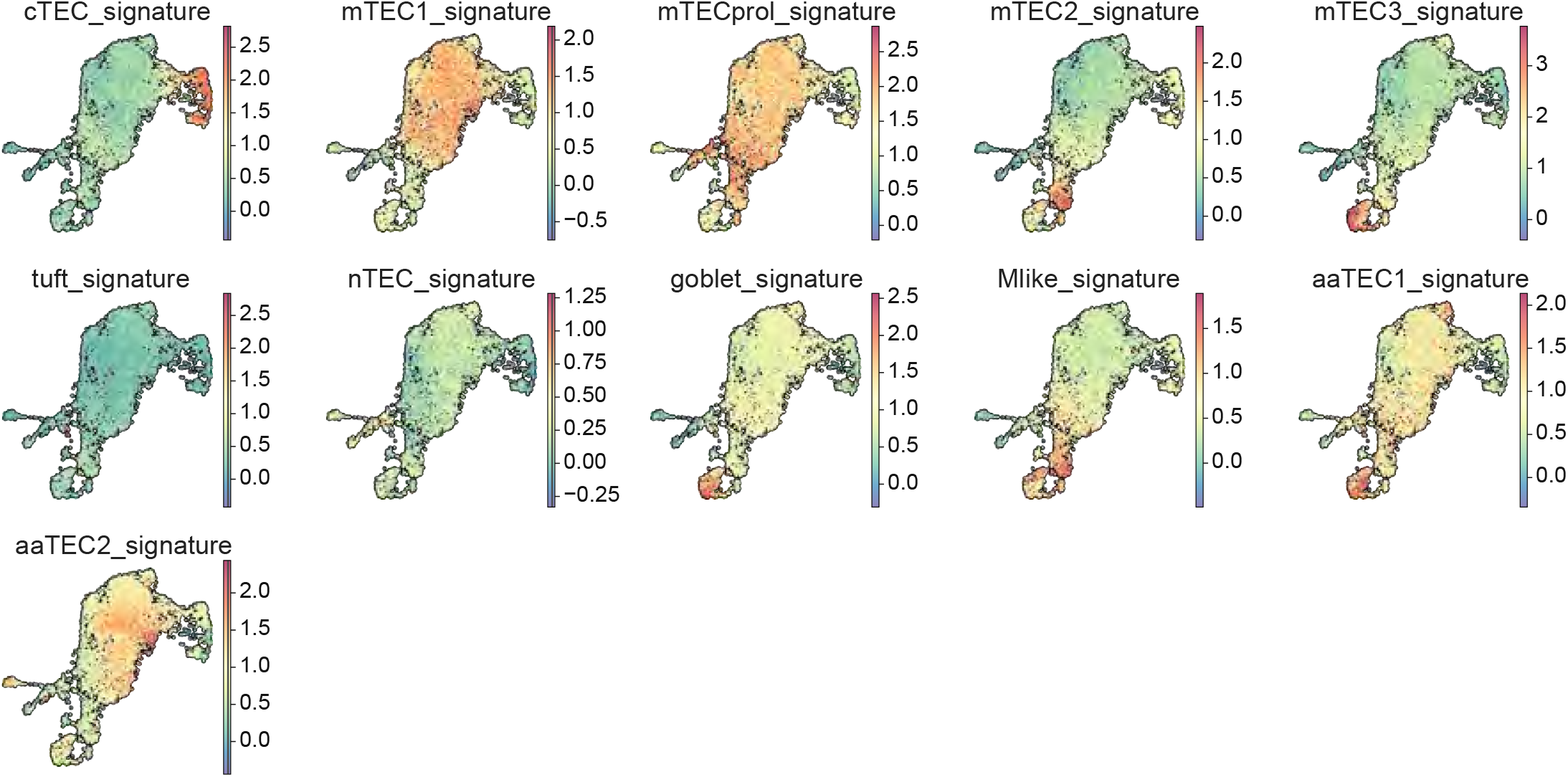
Signatures for epithelial cell subsets (**Data S3**), including aaTEC overlaid onto human TEC data derived from (*21*).

**Fig. S7:**
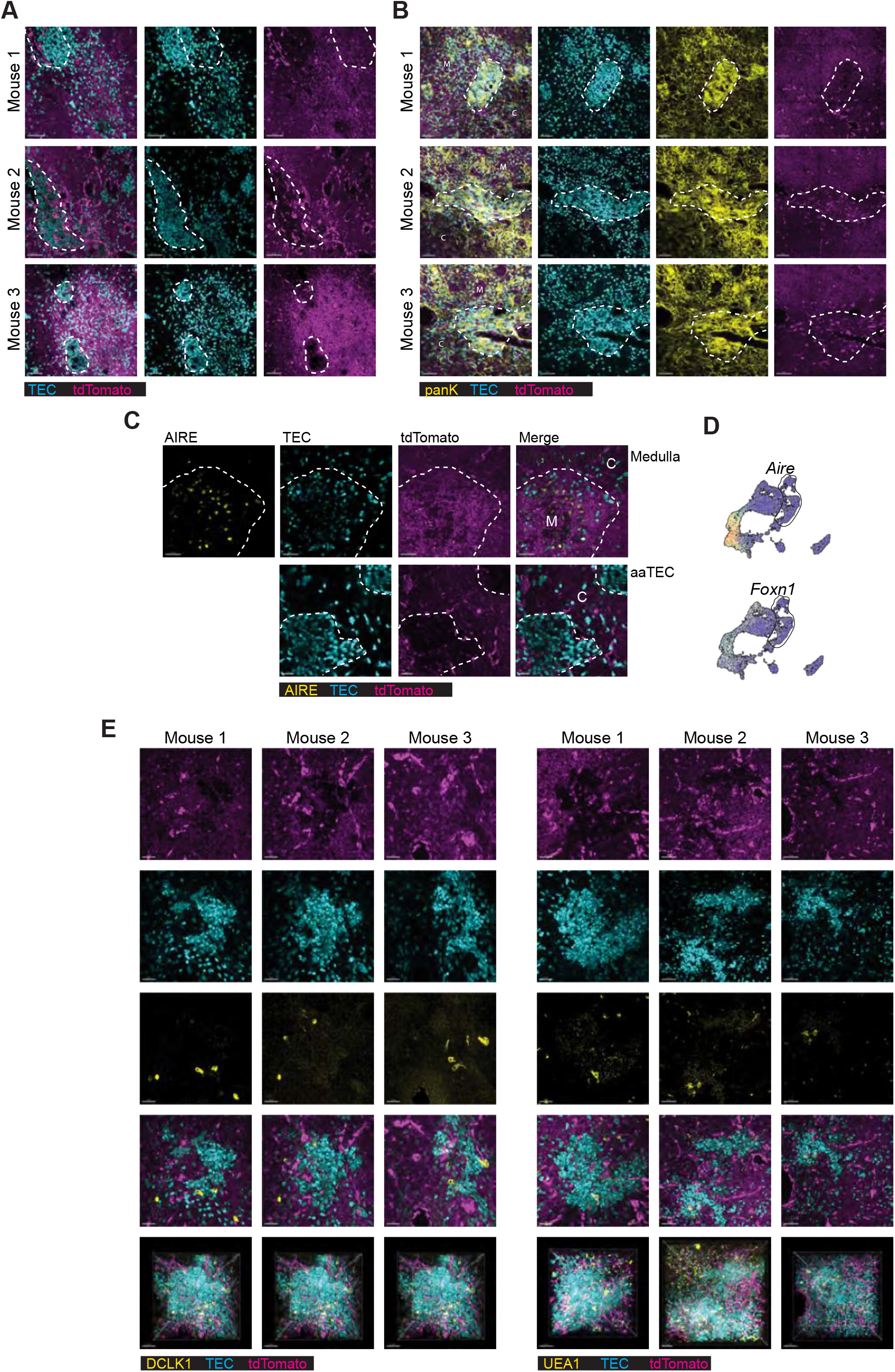
**A:** Representative confocal images of thymic sections from 12mo *Foxn1^nTnG^* mice with high-density TEC located in peri-medullary region. **B,** Representative confocal images of thymic sections from 12mo *Foxn1^nTnG^*mice stained with anti-pan-keratin, with high-density TEC regions highlighted. **C,** Representative confocal images of thymic sections from 12mo *Foxn1^nTnG^* mice stained with anti-AIRE, with the medulla or high-density TEC highlighted. **D**, *Aire* and *Foxn1* expression in TEC subsets. **E**, 3-D reconstruction and representative images of high-density TEC region from 12mo *Foxn1^nTnG^*mice stained with DCLK1 or UEA1 to highlight tuft cells and M-like cells, respectively. Scale bar: 50μm.

**Fig. S8:**
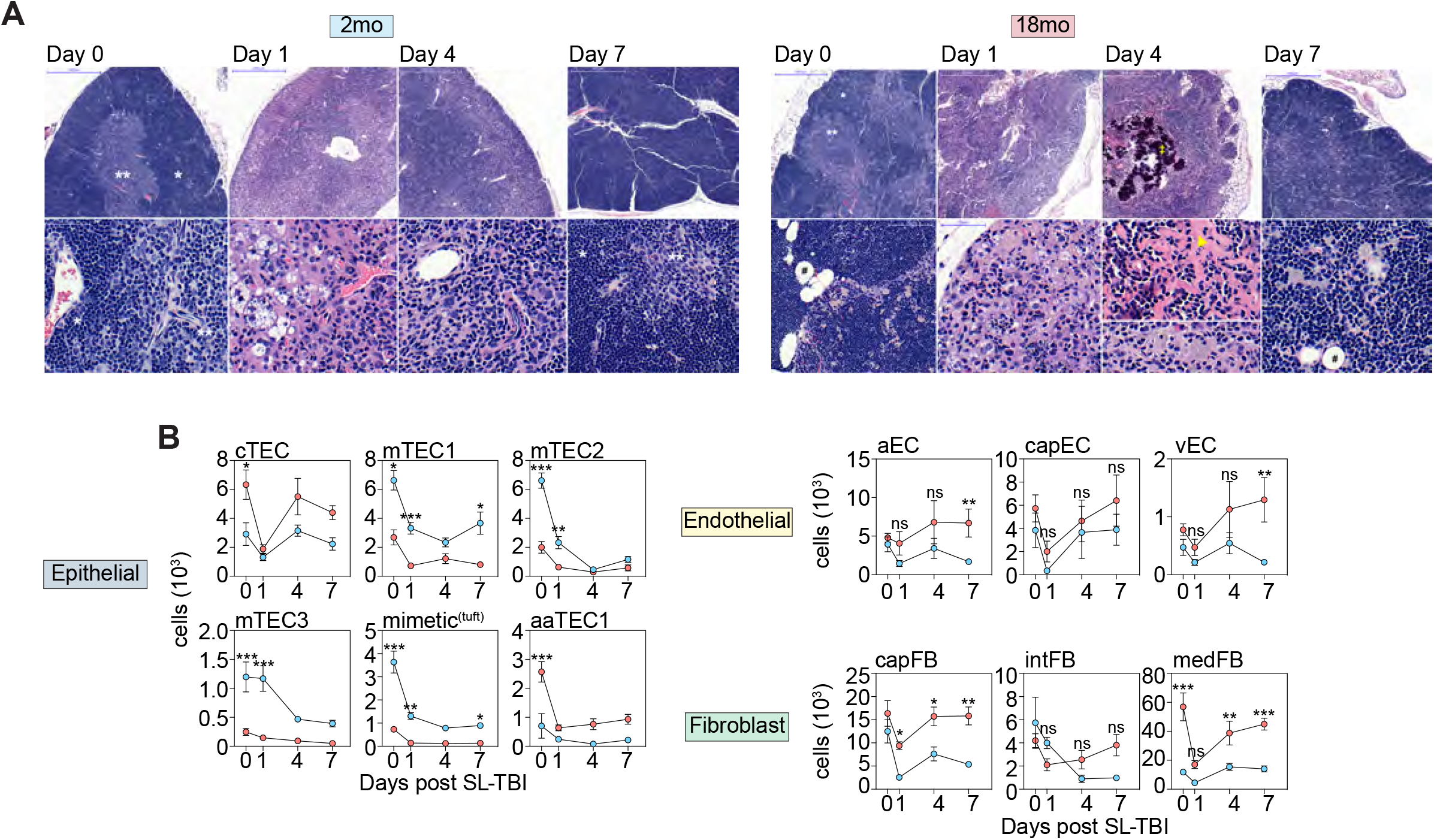
**A,** Morphologic alterations in the context of acute thymic involution after TBI and thymic reconstitution in 2mo and 18mo mice. Top and bottom rows represent low- and high-power images from each timepoint, respectively. Annotations: cortex (*), medulla (**), adipocytes (#), areas of dystrophic calcification (†), areas of dense fibrosis (arrowhead). **B-C**, Kinetics of recovery for the epithelial, endothelial and fibroblast defined subsets on day 0, 1, 4 and 7 after TBI in 2mo and 18mo mice. **B,** Total cellularity for each subset. Statistics compares across ages for each timepoint. Asterisks denote when recovery in either age cohort on a specific timepoint was significantly different to the other cohort. Summary data represents mean ± SEM; *, p<0.05; **, p<0.01; ***, p<0.001 using a two-way ANOVA.

**Fig. S9:**
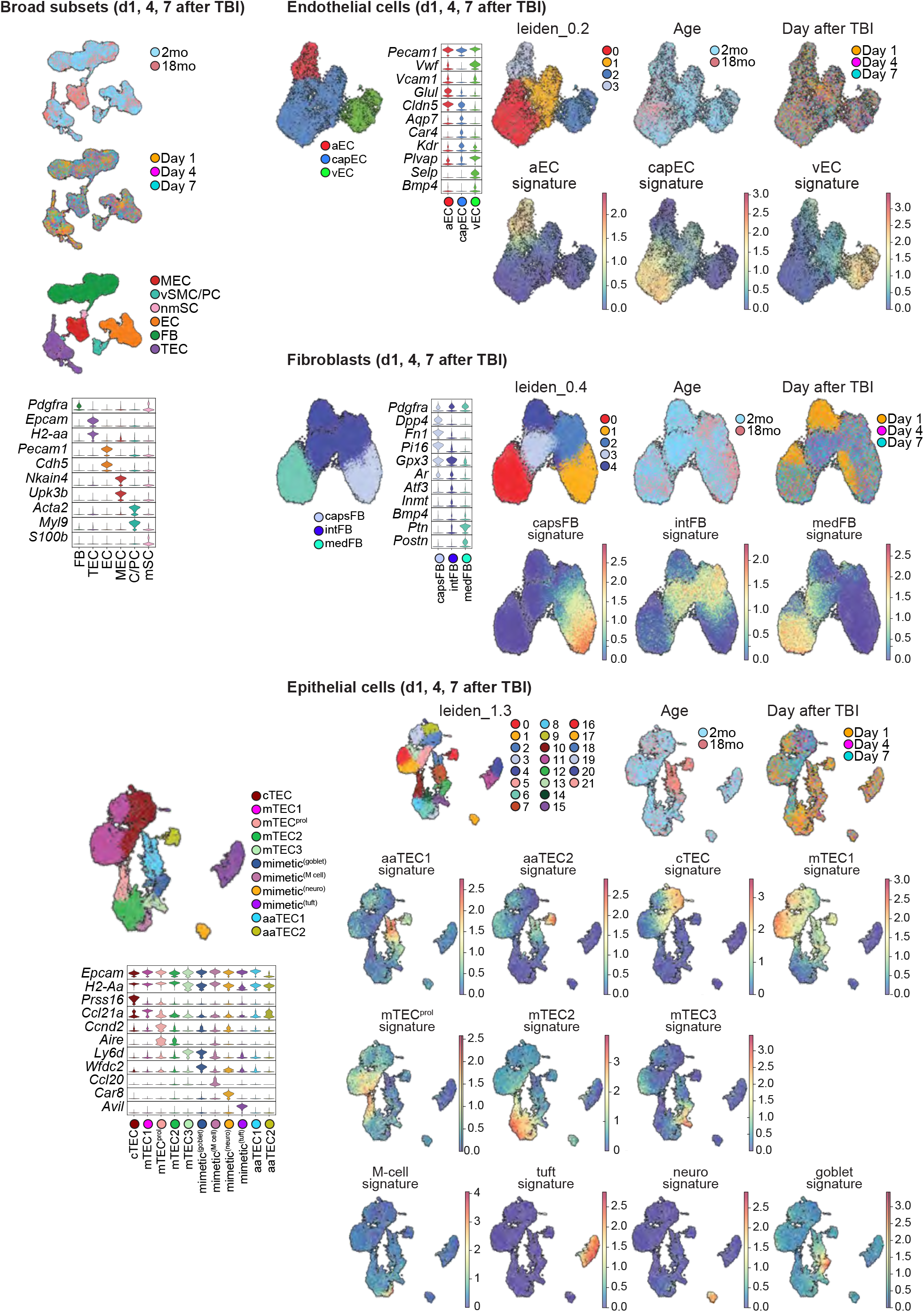
Leiden clustering and cell subset signatures derived from **Data S2** across endothelial, fibroblast and epithelial cells isolated at days 1, 4, and 7 after TBI.

**Fig. S10:**
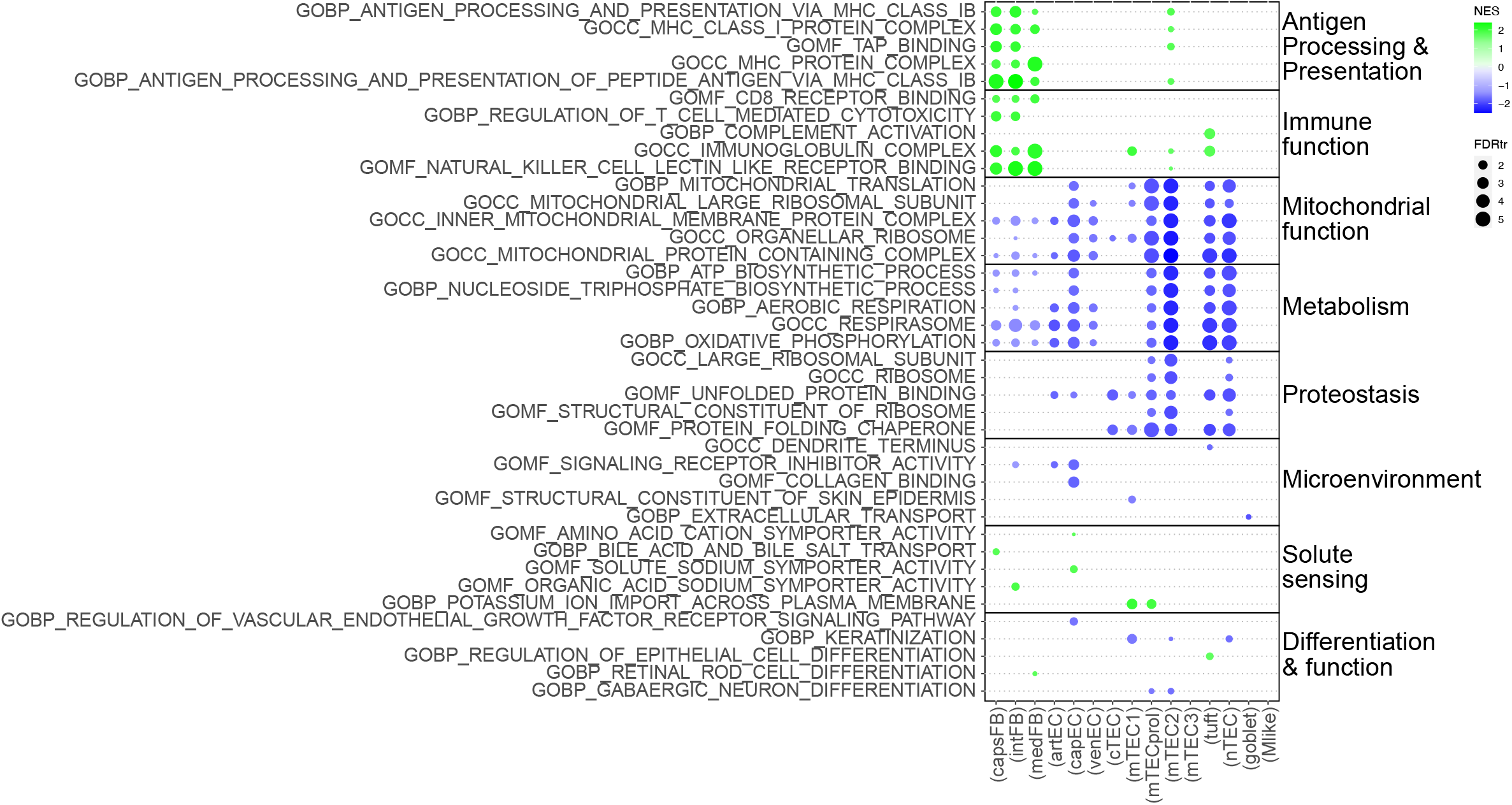
As in **Fig. 4A**, GSEA pathway analysis was performed for each subset based on differentially expressed genes within each population between 2mo and 18mo mice (**Data S4**). Dotplot of top 5 pathways within each category. Individual pathways are listed.

**Fig. S11:**
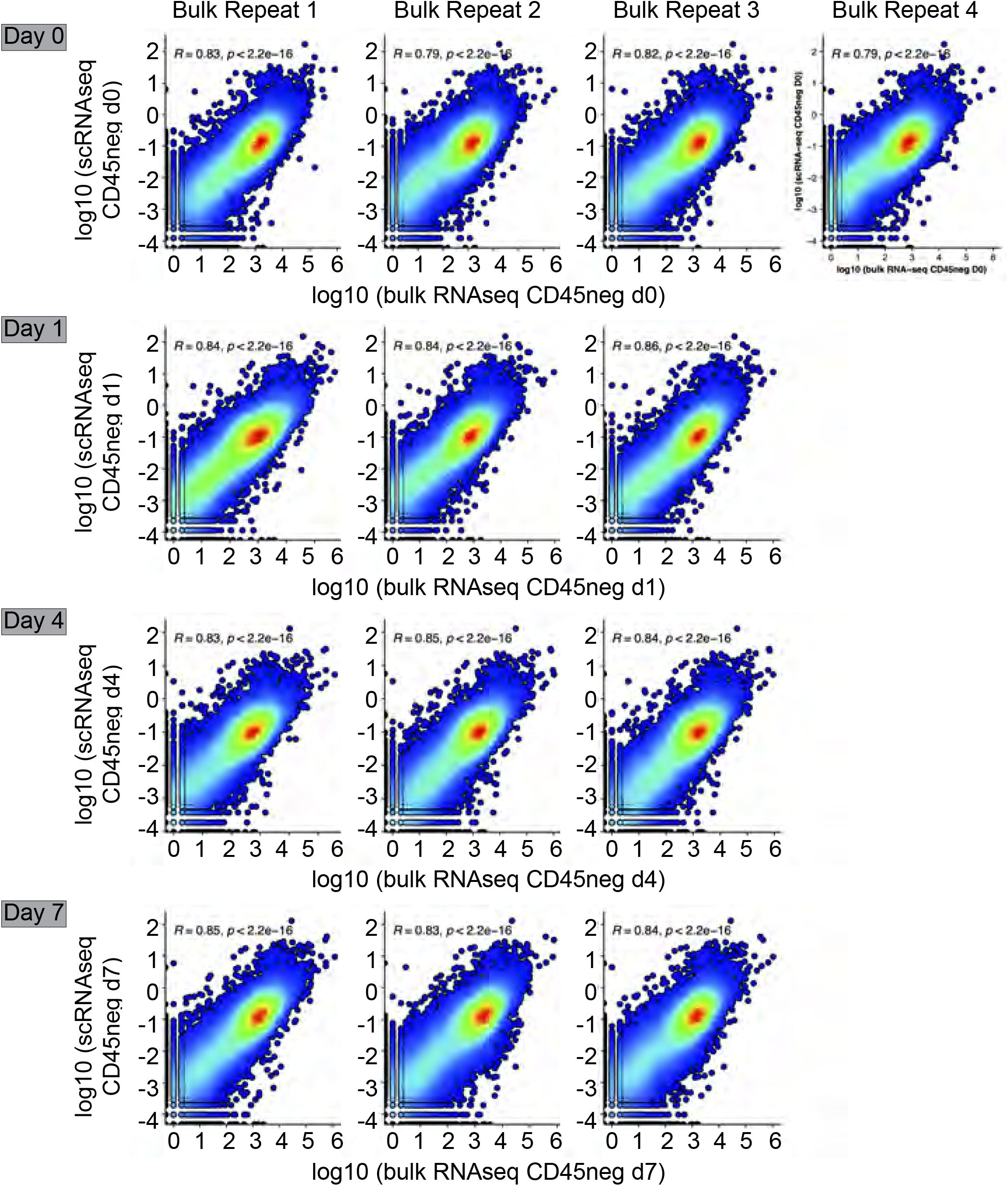
Scatterplots showing correlation (Pearson) between averaged expression profiles of CD45^-^ scRNA-seq averaged technical replicates collected at steady state (row 1), day 1 (row 2), day 4 (row 3) and day 7 (row 4) after TBI versus the expression profiles of CD45^-^ bulk RNA-seq biological replicates (replicate 1; column 1, replicate 2; column 2, replicate 3; column 3, replicate 4; column 4) obtained on the same timeframe.

**Data S1. Canonical markers for thymic stroma subsets.** Includes lists of canonical marker genes from thymic epithelial, endothelial and fibroblast defined subsets and how our thymic stroma subsets map to this published work.

**Data S2. Data Sof DEGs within thymic epithelium, endothelium and fibroblasts at steady-state**. Includes Wilcoxon rank metric results from all pairwise comparisons within the thymic epithelial, endothelial and fibroblast subsets at steady-state.

**Data S3. Data Sof top DEGs per subset used to create gene signatures**. Sheet 1 includes the top 30 DEGs per subset (sorted by Wilcoxon rank score in decreasing order), and their human orthologues used to map our epithelial, endothelial and fibroblast subsets to the human CD45^-^ dataset by Park et. al. Sheet 1 includes the top 10 DEGs per subset (sorted by Wilcoxon rank score in decreasing order) used to create subset-specific gene signatures to map our subsets to the populations found at days 1, 4 and 7 after TBI.

**Data S4. GSEA enriched pathways with aging.** Includes EnrichmentMap-generated network Data Swith the combined GSEA results from all 18mo vs 2mo steady-state subset comparisons. To assist result interpretation, pathways have been manually categorized and labeled based on gene overlap and pathway relevance in respect to Aging Hallmarks.

**Data S5. GSEA enriched pathways for mTEC1 and aaTECs.** Includes GSEA results for mTEC1 (comparison: 18mo vs 2mo) and aaTEC1/2 (comparison: aaTEC1/2 vs TECs). Individually enriched pathways shown in **Fig. 2J** including the genes contributing to their enrichment are also listed here. GOBP_KERATINIZATION (mTEC1), GOBP_AP&P_OF_PEPTIDE_ANTIGEN (aaTEC1); HALLMARK_EMT (aaTEC2).

**Movie S1.**

Associated with Fig. 2B. Whole tissue light sheet imaging showing central medulla (magenta) in 2mo (A) and 18mo (B) female *Foxn1^nTnG^* mice with HD-TEC (cyan) only apparent in old mice.

**Movie S2.**

Associated with Fig. 3E. Whole tissue imaging of 12mo Foxn1^nTnG^ mice at baseline (A) or 28 days after TBI (B).

## METHODS

### Isolation of cells and flow cytometry

All steps were performed at 4 C unless indicated. 10-15 thymi of 2-month-old or 18-month-old female C57BL/6 mice were excised and enzymatically digested following and adapted protocol (*77, 78*). Briefly, thymi were mechanically dissociated into 2 mm pieces. Tissue pieces were incubated with a digestion buffer (RPMI, 10% FCS, 62.5 um/mL liberase TM, 0.4 mg/ml DNase I) twice for 30 min at 37 C. Between incubation steps, supernatant containing dissociated cells was transferred to 50 mL conical tubes equipped with 100 um filter. Cells were pelleted by centrifugation at 400g for 5 min. For sequencing experiments, cell pellets were incubated with anti-mouse CD45 microbeads and CD45^+^ cells were depleted from cell suspension using magnetic-associated cell sorting (MACS) on LS columns according to manufacturer’s protocol. Following red blood cell lysis using ACK buffer, the CD45-depleted cell fraction was incubated with an antibody cocktail for 15 min at 4 C and cells of interest were purified by fluorescent-associated cell sorting (FACS) on a BD Biosciences Aria II using a 100 um nozzle. Cells were sorted into tubes containing RPMI supplemented with 2% BSA. FACS-purified cells were spun down at 400g for 5 min and resuspended in PBS supplemented with 0.04 % BSA for generation of single-cell suspensions.

For flow cytometry and cell sorting, surface antibodies against CD45 (30-F11), CD31 (390 or MEC13.3), TER-119 (TER-119), MHC-II IA/IE (M5/114.15.2), EpCAM (G8.8), Ly51 (6C3), PDGFRα (APA5), CD104 (346-11A), L1CAM (555), Ly6D (49-H4), Gp38 (8.1.1), CD26 (H194-112), CD62P (RB40.34), and CD309 (Avas12a) were purchased from BD Biosciences (Franklin Lakes, NJ), BioLegend (San Diego, CA) or eBioscience (San Diego, CA). Ulex europaeus agglutinin 1 (UEA1), conjugated to FITC or Biotin, was purchased from Vector Laboratories (Burlingame, CA). Flow cytometry was performed on a Fortessa X50 (BD Biosciences, Franklin Lakes, NJ) and cells were sorted on an Aria II (BD Biosciences) using FACSDiva (BD Biosciences, Franklin Lakes, NJ). Analysis was performed by FlowJo (Treestar Software, Ashland, OR).

### Cell Preparation for sequencing

The single-cell RNA-seq of FACS-sorted cell suspensions was performed on Chromium instrument (10X genomics) following the user guide manual (CG00052 Rev E) and using Single Cell 3’ Reagent Kit (v2). The viability of cells prior to loading onto the encapsulation chip was 73-98%, as confirmed with 0.2% (w/v) Trypan Blue stain. Each sample, containing approximately 8000 cells, was encapsulated in microfluidic droplets at a final dilution of 66–70 cells/µl (a multiplet rate ∼3.9%). Following reverse transcription step the emulsion droplets were broken, barcoded-cDNA purified with DynaBeads and amplified by 12-cycles of PCR: 98 °C for 180 s, 12x (98 °C for 15 s, 67 °C for 20 s, 72 °C for 60 s), and 72 °C for 60 s. The 50 ng of PCR-amplified barcoded-cDNA was fragmented with the reagents provided in the kit, purified with SPRI beads and resulting DNA library was ligated to the sequencing adapter followed by indexing PCR: 98 °C for 45 s; 12x 98 °C for 20 s, 54 °C for 30 s, 72 °C for 20 s), and 72 °C for 60 s. The final DNA library was double-size purified (0.6–0.8X) with SPRI beads and sequenced on Illumina Nova-Seq platform (R1 – 26 cycles, i7 – 8 cycles, R2 – 70 cycles or higher) at depth of 200-270 million reads per sample.

### CD45^-^ single cell RNA-seq preprocessing and downstream data analysis

FASTQ files were processed using the 10x Cell Ranger package (v7.01). The Cell Ranger generated filtered_feature_bc_matrix.h5 files were processed following the guidelines on the shunPykeR GitHub repository (*79*), an assembled pipeline of publicly available single cell analysis packages put in coherent order, that allows for data analysis in a reproducible manner and seamless usage of Python and R code. Genes that were not expressed in any cell and also ribosomal and hemoglobin genes were removed from downstream analysis. Each cell was then normalized to a total library size of 10,000 reads and gene counts were *log*-transformed using the *log*(X+1) formula, in which *log* denotes the natural logarithm. Principal component analysis was applied to reduce noise prior to data clustering. To select the optimal number of principal components to retain for each dataset, the knee point (eigenvalues smaller radius of curvature) was used. Leiden clustering (*79*) was used to identify clusters within the PCA-reduced data.

Quality of the single cells was computationally assessed based on total counts, number of genes, mitochondrial and ribosomal fraction per cell, with low total counts, low number of genes (≤1000) and high mitochondrial content (≥0.2) as negative indicators of cell quality. Cells characterized by more than one negative indicator were considered as “bad” quality cells. Although cells were negatively sorted prior to sequencing for the CD45 marker, a small amount of CD45^+^ cells (expressing *Ptprc*), and also a few parathyroid cells (expressing *Gcm2*), were detected within our dataset. To remove bad quality cells and contaminants in an unbiased way, we assessed them in a cluster basis rather than individually. Leiden clusters with a “bad” quality profile and/or a high number of contaminating cells were removed. Finally, cells marked as doublets by scrublet (*80*) were also filtered out. Overall, a total of 12,497 cells, representing 13.3% of all our data, was excluded from further analysis (see Figure S2 for per sample metrics). After removal of these cells, PCA and unsupervised clustering analysis was reapplied to the filtered data as described above, followed by batch effect correction across all samples using harmony (*81*).

The shunPykeR adapted jupyter notebooks to replicate preprocessing, downstream analysis and figure reproducibility of this manuscript can be found at <u>github.com/kousaa/KousaJahnZhao_et_al_2023</u>.

#### Differential expression analysis

Differential expression analysis for comparisons of interest was performed using the Wilcoxon test (*82*). In all cases, differentially expressed genes were considered statistically significant if the FDR-adjusted *p*-value was less than 0.05.

#### Generation of cell type/subset-specific gene signatures

To select maximally specific genes per cell type/subset, we ran MAST for all pairwise cluster comparisons (each versus the rest) and retained the top 10 DEGs for a given cluster (FDR<=0.05, sorted by decreasing coef). To generate the unique cluster signatures per se, we used the *sc.tl.score_genes()* function from *scanpy* (*83*) that calculates averaged scores based on the cluster specific genes (scores are subtracted with a randomly sampled reference gene set).

#### Mapping steady-state subsets onto the TBI timeseries

To identify the steady-state structural cell types and subsets within the day 1, 4 and 7 after TBI, we first subsetted and reanalyzed our datasets for only the days after damage. Due to the huge transcriptional shift that happens on day 1 after TBI, we calculated highly variable genes excluding samples from day 1, and then applied batch effect correction across all samples with harmony. We then applied unsupervised clustering using leiden and assigned cell identities using (a) canonical markers for the major cell types, (b) steady-state gene signatures from the steady-state subsets (top 10 DEGs, wilcoxon: FDR<=0.05 & sorted in descending order by score, **Fig. S9** and **Data S2**). Once annotated, we combined the 2mo and 18mo time series datasets in one UMAP (**Fig. 3H**).

#### ThymoSight

ThymoSight is an R Shiny app we have developed to allow interactive exploration of single cell datasets of the non-hematopoietic thymic stroma. The app.R code that launches the app together with the python notebook used to create consistent annotation fields, reanalyze and integrate the public datasets with ours have been submitted on GitHub (https://github.com/kousaa/ThymoSight). The server hosting the interactive app can be accessed at www.thymosight.org.

#### Pathway enrichment analysis

Pathway enrichment analysis was performed with GSEA (Subramanian et al., 2005) according to the gene list and rank metric provided. The GSEA Preranked module was used to predict pathway enrichment in threshold free comparisons: (a) 18mo vs 2mo subsets at steady-state and (b) aaTEC1 and aaTEC2 versus other TEC. We created rankings for all differentially expressed genes using the Wilcoxon Z-score in descending order. Predicted pathways with an FDR<=0.05 were considered as significantly enriched.

#### Network analysis

Network analysis of the significantly enriched GSEA pathways from comparisons of interest was performed using the Cytoscape (*55*) module EnrichmentMap(Merico et al., 2010). We used EnrichmentMap to organize enriched pathways (FDR<=0.05) with a high overlap of genes (default cutoff similarity: 0.375) in the same network allowing for a simplified and intuitive visualization of the distinct processes that are significantly represented in the subset/stage and day transitions of interest. This facilitated interpretation of the enriched pathways from the plethora of comparisons and allowed categorization of all resulting pathways into networks based on the overlap of the genes contributing to the pathway’s enrichment. Manual inspection of the resulting networks allowed allocation of network-related annotations. Individual pathways that were not part of an existing network were manually annotated to the existing categories based on their biological function or grouped under “other”.

### CD45^-^ bulk RNA-seq preprocessing and downstream data analysis

#### Quality control, alignment and gene count quantification

Quality control (QC) of the raw read files (FASTQ) was performed using the FastQC tool. Low quality reads and adapter contaminants were removed using Trimmomatic (*84*) (default parameters for paired-end reads) and post-trimmed reads were re-assessed with FastQC to verify adapters removal and potential bias introduced by trimming. QC-approved reads were aligned to the GRCm38.p5 (mm10) mouse genome assembly (GENCODE; M12 release) with STAR (*85*) aligner using default parameters and *--runThreadN* set to 32 to increase execution speed. The STAR-aligned files were then used as input to the featureCounts (*86*). tool (default parameters) to quantify gene expression levels and construct the count matrix.

#### Low gene count removal and library size normalization

The raw count matrix was converted to a DGEList object in R using the *readDGE()* function from the edgeR (*87*) package. Lowly expressed genes were removed using the *filterByExpr()* function for the groups of interest prior to comparison with the scRNA-seq datasets.

#### Bulk RNA-seq vs scRNA-seq

scRNA-seq sample reproducibility was verified using bulk RNA-seq data for the CD45^-^ sorted populations. Comparison between bulk and single cell RNA-seq CD45^-^ transcriptional profiles was performed by computing Pearson’s correlation between *log*_10_- transformed raw bulk counts (per biological replicate) and log_10_-transformed averaged raw single cell counts (per technical replicate) for the relevant datasets across the TBI timeframe (**Fig. S11**).

### Thymic tissue clearing and immunofluorescence

After euthanasia, mice were transcardially perfused with PBS followed by 4% PFA. Thymi were dissected and post-fixed in 4% PFA for 4h at 4°C. For confocal imaging, fixed tissue was sectioned at 200um using a Leica VT1000 S vibratome. Tissue clearing was performed as previously described (Chung et al., 2013, p. 201) with some modifications. Briefly, tissue was immersed in monomer buffer (4% acrylamide, 0.25% (w/v) azo-initiator (Wako Pure Chemical Industries Ltd., Osaka, Japan) in PBS) and incubated at 4°C overnight. The solution was transferred to a vacuum tube and bubbled with nitrogen gas for 15min. The gel was set for 3h at 37°C with gentle rotation after which the tissue was transferred to clearing buffer (8% SDS and 50mM sodium sulphite in PBS) and cleared at 37°C until turning semi-transparent. To remove SDS, samples were transferred to the following buffers to wash for 1h each with rotation: 1) 1% SDS, 0.5% Triton-X in PBS; 2) wash buffer (1% BSA and 0.5% Triton-X in PBS), 2 washes; 3) blocking buffer (4% normal serum, 1% BSA and 0.3% Triton-X in PBS), 2 washes. Antibodies were diluted in blocking buffer at the dilutions indicated below. The antibodies used were rabbit anti-pan-Cytokeratin (Dako, Cat# Z0622), anti-K5 (BioLegend, Cat# poly19055), rat anti-mouse K8/18 (Troma-1; Developmental Studies Hybridoma Bank, Iowa City), rabbit anti-K14 (AbCam, Cat# EPR17350), rat anti-mouse AIRE (WEHI, Clone# 5H12), rabbit anti-human/mouse DCLK1 (LSBio, Cat# LS-C100746), biotinylated UEA-1 lectin (Vector labs, USA, Cat# B-1065). The secondary antibodies used were Alexa Fluor® 647 Donkey anti-rabbit IgG (H+L) (Invitrogen, Cat# A31573), Alexa Fluor® 647 Goat anti-rat IgG (H+L) (Invitrogen, Cat# A-21247), Alexa Fluor™ 647 Streptavidin conjugate (Invitrogen, Cat# S21374). After staining, samples were washed in PBS with 0.3% Triton-X. For imaging, samples were incubated in EasyIndex optical clearing solution (RI=1.46) (LifeCanvas Technology) at room temperature until turning fully transparent.

#### Confocal microscopy imaging

Tissue sections were imaged on a Zeiss LSM 880 confocal microscope using a Plan-Apochromat 25x/0.8 multi-immersion objective at a voxel size of 0.22um in XY and 2um in Z.

#### Light sheet microscopy imaging

Whole thymic lobes were scanned using a Zeiss Z.1 Lightsheet microscope. The detection objective was an EC Plan-Neofluar 5x/0.16. Stacks were acquired at a resolution of 0.915 um in XY and approximately 4.9 um in Z. Dual-side images were fused using the maximum intensity option.

#### Image presentation

All images shown are processed using Imaris 9.7.1 (Bitplane). Regions of interest in tissue sections are presented as 10um Z-projections.

#### Volume calculation of thymic regions

The total volumes of entire right thymic lobes were calculated using Imaris by generating the lobe surface from the tdTomato channel. Medullary regions were defined by high GFP^+^ cell density and Krt14^+^, and High-density (HD-TEC) region in aged thymus was defined by compacted GFP^+^ TECs. To identify medullary and HD TEC regions in the images, we developed a pipeline in ImageJ (Version: 2.3.0/1.53f) (*88*). For medulla, GFP and Keratin 14 channel were combined. For HD TEC region, only GFP^+^ channel was used. The images were filtered using 2D median and 3D Gaussian filtering and then binarized using a 2D min/max filter with thresholds set according to fluorescence intensity. The resulting image was used to extract the medullary or HD TEC surface. The cortical volume was calculated as total thymic lobe volume minus the medullary and HD TEC volume.

#### Segmentation of TEC nuclei

TEC nuclei were identified in confocal images of thymus sections using the spot detection function in Imaris. Total TEC spots were filtered and then TEC subsets were segmented according to the shortest distance to the indicated surface or the section edge. Medullary or HD TECs were defined by the shortest distance to indicated surface <= 0 um, sub-capsular TEC were defined as shortest distance to section edge >= -25 um and the remaining spots were defined as cTECs.

Mean cell density was calculated by dividing numbers of specific TEC subset to the volume of different thymic region.

#### Quantification of TECs

The number of nuclei in the various TEC subsets in the right lobe was calculated by multiplying the mean cell densities ascertained by confocal analysis of the slices from the left lobe by the volumes determined by light sheet imaging of the right lobe.

### Statistics

Statistical analysis between two groups was performed with the nonparametric, unpaired Mann-Whitney U test. Statistical comparison between 3 or more groups was performed with the nonparametric, unpaired Kruskall-Wallis test. All statistics were calculated and display graphs were generated in Graphpad Prism.

## Notes

### Summary of Updates

In this revised manuscript we have used a variety of advanced approaches to interrogate the emergence of age-associated TECs (aaTECs) and their function during age and damage, including spatial transcriptomics, genetic lineage-tracing, whole organ light sheet and confocal imaging, and advanced bioinformatic approaches. Notably, this expanded scope of the manuscript involves new collaborations, including inclusion of a new senior author, Dr. Daniel Gray, and new co-first author Kelin Zhao, from the Walter and Eliza Hall Institute. Their team has brought advanced imaging approaches, applied for the first time in the thymus, to show that these aaTECs form unusual high-density structures in the aged thymus and that after damage these aaTEC clusters rapidly expand after damage. Importantly, despite the loss of canonical TEC markers and markers of epithelial identity, we now definitively show with two lineage-tracing approaches by scRNAseq, flow cytometry, and imaging that aaTECs are derived from Foxn1+ TECs, but that these aaTEC regions do not support thymocyte development. Our findings suggest that aaTECs limit thymic function by: 1) replacing functional thymic tissue with high density 'scars', and 2) drawing growth signals such as BMP and FGF signaling from conventional TECs. Furthermore thymic regeneration following damage was diminished with age coincident with the expansion of aaTEC, with an overall reduction in key growth factors. Finally, we provide evidence that inflammaging signals and senescence-associate secretory phenotype in thymic fibroblasts drive the emergence of aaTECs. It is also notable that many of these factors have also been implicated in driving EMT.

